# Temperature-dependent gene regulatory divergence underlies local adaptation with gene flow in the Atlantic silverside

**DOI:** 10.1101/2022.04.27.489786

**Authors:** Arne Jacobs, Jonathan P. Velotta, Anna Tigano, Aryn P. Wilder, Hannes Baumann, Nina O. Therkildsen

## Abstract

Gene regulatory divergence is thought to play an important role in adaptation, yet its extent and underlying mechanisms remain largely elusive under scenarios of local adaptation with gene flow. Local adaptation is widespread in marine species despite generally high connectivity and often associated with tightly-linked genomic architectures, such as chromosomal inversions. To investigate gene regulatory evolution under gene flow and the role of discrete genomic regions associated with local adaptation to a steep thermal gradient, we generated RNA-seq data from Atlantic silversides (*Menidia menidia*) from two locally adapted populations and their F1 hybrids, reared under two different temperatures. We found substantial divergence in gene expression and thermal plasticity, with up to 31% of genes being differentially expressed, and primarily *trans*-rather than *cis*-regulatory divergence between populations, despite ongoing gene flow. Substantially reduced thermal plasticity, temperature-dependent gene misexpression and the disruption of co-expression networks in hybrids point toward a role of regulatory incompatibilities in maintaining local adaptation, particularly under colder temperatures, which appear more challenging for this species. Adaptive chromosomal inversions seem to play an important role in gene regulatory divergence through the accumulation of regulatory incompatibilities but are not consistently enriched for divergently regulated genes. Together, these results highlight that gene regulation can diverge substantially among populations connected by strong gene flow in marine environments, partly due to the accumulation of temperature-dependent regulatory incompatibilities within inversions.

## Introduction

Despite the lack of obvious barriers to gene flow in the oceans, it has become clear that local adaptation is widespread in marine species (Conover et al. 2006; Sanford and Kelly 2011; Kelley et al. 2016; Hays et al. 2021). There have been substantial advances in our understanding of the genomic basis and architecture of local adaptation with gene flow in recent years, with many studies identifying large divergent haplotype blocks often linked to chromosomal inversions that appear to play a key role (Kirubakaran et al. 2016; Sodeland et al. 2016; Pettersson et al. 2019; Han et al. 2020; Wilder et al. 2020; Tigano et al. 2021). However, we are still lacking a detailed understanding of the molecular mechanisms associated with local adaptation with gene flow. While gene regulatory variation has been argued to be a major target of selection during local adaptation (Pavey et al. 2010; Mack and Nachman 2017; McGirr and Martin 2020a), the regulatory mechanisms underlying divergence in gene expression under gene flow and how this is associated with the often tightly-linked genomic architecture of divergence (e.g. through inversions), remains largely unknown.

In contrast to sequence variation, gene expression is not affected by recombination, and the analysis of gene expression therefore offers important opportunities to dissect the mechanisms of local adaptation and identify adaptive genes despite the presence of large inversion polymorphisms that suppress recombination across large genomic blocks (Said et al. 2018; Crow et al. 2020; Berdan et al. 2021). In general, adaptive divergence within a species is thought to be facilitated by *trans*-regulatory changes, such as variation in transcription factors (Hart et al. 2018; McGirr and Martin 2020b), as *trans*-changes likely affect many downstream genes in regulatory networks, have a larger mutational target size and are more likely to persist as polymorphisms within species (Mack and Nachman 2017; Cutter and Bundus 2020). However, under strong gene flow and the presence of tightly-linked genomic architectures (e.g. inversions), more modular and tissue-specific *cis*-acting changes might be advantageous, as co-evolved factors (e.g. *cis*-regulatory elements and target gene(s)) remain linked and are less likely to be broken up by homogenising gene flow (Wittkopp et al. 2004; Wittkopp and Kalay 2011; Gould et al. 2018; Said et al. 2018; Verta and Jones 2019; Crow et al. 2020; Berdan et al. 2021). On the other hand, inversions can globally affect gene expression by harbouring divergent *trans*-factors (Naseeb et al. 2016; Said et al. 2018). Thus, even if genome scans suggest that population divergence is concentrated within narrow genomic regions in many cases of local adaptation with gene flow, *trans*-effects of inversions could drive changes in functionally divergent genes spread throughout the genome.

*Cis*-and *trans*-regulatory changes are not mutually exclusive and co-evolve to reach or maintain optimal adaptive expression levels (Fraser et al. 2010). Thus, co-evolution of *cis*-and *trans*-regulatory variation can lead to the emergence of Dobzhansky-Muller hybrid incompatibilities between populations (Mack and Nachman 2017; Signor and Nuzhdin 2018; McGirr and Martin 2020a). The maintenance of local adaptation despite gene flow might suggest the evolution of such regulatory incompatibilities between populations (Ortíz-Barrientos et al. 2007; Mack and Nachman 2017), which can manifest in ‘hybrid gene misexpression’, i.e. the transgressive expression of genes, that is expression levels in hybrids are higher or lower than their parental expression range (Landry et al. 2005; Landry et al. 2007; Ortíz-Barrientos et al. 2007; Mack and Nachman 2017). Hybrid gene misexpression has been observed in some cases in the early stages of speciation, suggesting that regulatory incompatibilities can arise early in the divergence process (Renaut et al. 2009; Barreto et al. 2015; McGirr and Martin 2020a). In particular, incompatible regulatory alleles might accumulate between non-recombining inversion-linked haplotypes (Berdan et al. 2021). However, gene regulatory mechanisms can be highly context-dependent, meaning that gene regulatory variation and/or incompatibilities, and hence inversions, might only act and become visible in specific environments, tissues, sexes and/or life-stages (York et al. 2018; Mugal et al. 2020; Berdan et al. 2021; Hu et al. 2022). It remains unclear, if such regulatory incompatibilities and associated hybrid misexpression can arise under ongoing pervasive gene flow, and if so, how this is facilitated by tightly-linked genomic architectures.

In this study, we investigated the gene regulatory architectures and mechanisms underlying local adaptation in a highly-connected marine teleost, the Atlantic silverside (*Menidia menidia*). Atlantic silversides are distributed along one of the steepest temperature clines in the world along the North American Atlantic Coast (Hice et al. 2012) and show a high degree of local adaptation across their range (Conover and Heins 1987; Yamahira and Conover 2002; Hice et al. 2012) despite high gene flow (Mach et al. 2011; Lou et al. 2018; Wilder et al. 2020). Countergradient variation in growth rate, with intrinsic growth rates generally higher in Northern latitudes to compensate for shorter growing seasons, is a major adaptation to stark differences in thermal regimes, as are genetically based differences in many other traits, e.g. fecundity, metabolic rate, swimming performance, and foraging behavior (Yamahira and Conover 2002; Conover et al. 2009; Hice et al. 2012). Multiple large inversions (0.4 to 12.5 Mb) display fixed differences for opposite orientations (co-linear vs. inverted) across the latitudinal cline (Wilder et al. 2020; Tigano et al. 2021; Akopyan et al. 2022), with otherwise minimal genetic background differentiation between populations, suggesting that chromosomal rearrangements that locally reduce recombination (Akopyan et al. 2022) play a key role in maintaining co-adapted alleles despite gene flow.

Given the strong phenotypic divergence in the face of gene flow (Conover et al. 2009; Hice et al. 2012), we hypothesise that gene expression and temperature-responses are strongly divergent between locally adapted Atlantic silverside populations, and that divergently regulated genes are concentrated within the large inversions that show highly elevated levels of differentiation between populations. Furthermore, we expect that *cis*-regulatory differences are predominant due to strong linkage among co-adapted alleles, particularly within inversions, which potentially leads to the presence of misexpression in hybrids due to incompatible regulatory variation. To test these predictions, we analysed the genome-wide gene expression in Atlantic silversides from locally-adapted populations, and their F1 hybrids, reared under two temperatures resembling their local and non-local environments, respectively, during their growth phase (Fig. 1). By integrating analyses of gene (co-)expression within and across populations and temperatures, with gene expression inheritance and allele-specific expression patterns, we reveal strong temperature-dependent gene regulatory divergence despite gene flow. We furthermore show that divergently expressed genes are widespread across the genome and only partially enriched within some inversions, although inversions seem to harbour genetic incompatibilities leading to gene misexpression in heterokaryotypes. Overall, our results support a key role for gene regulatory divergence in enabling local adaptation with gene flow, the dependence of regulatory mechanisms on environmental conditions, and a highly context-specific role of inversions in driving gene expression differences.

**Figure 1.**
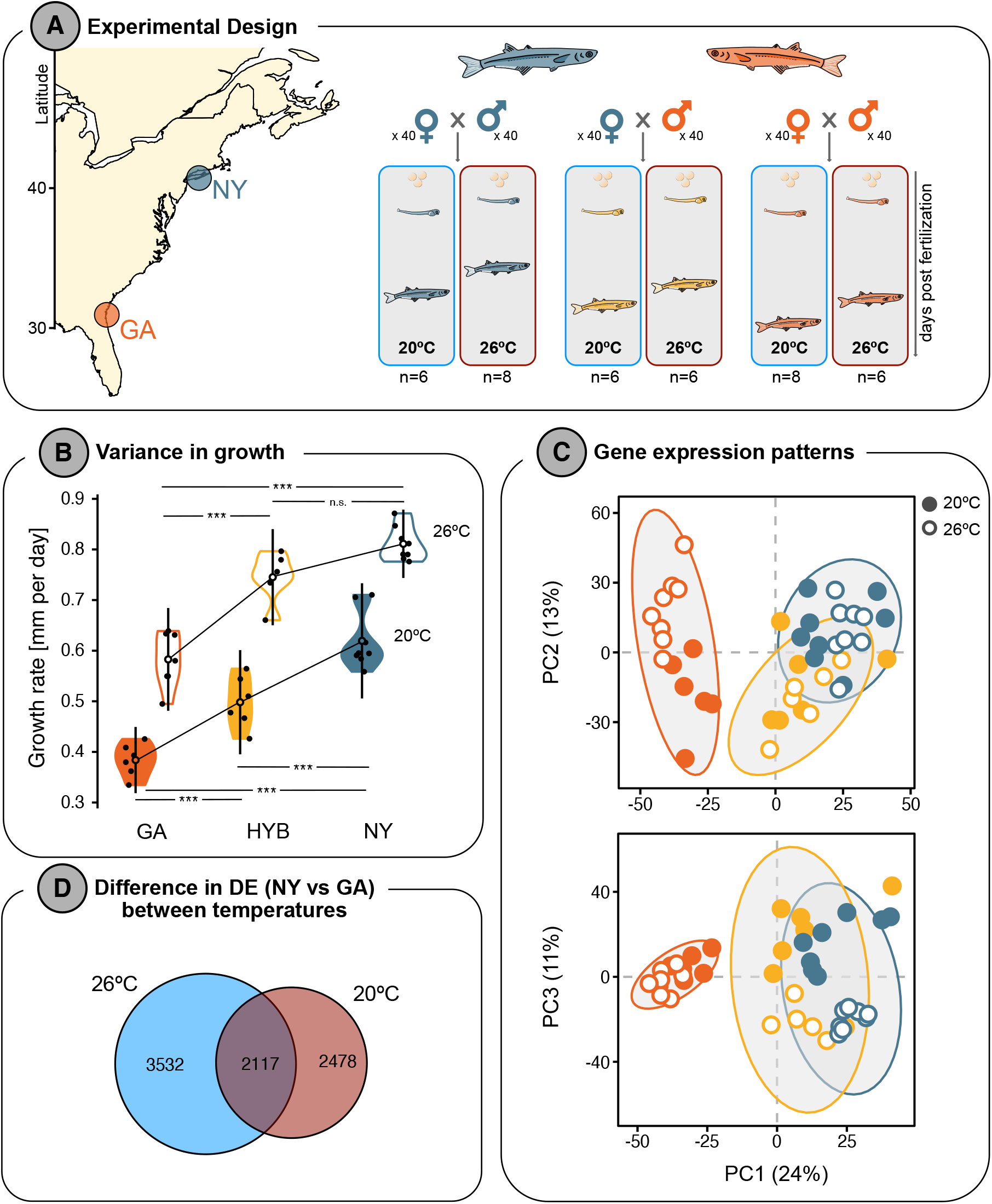
**A)** Common garden experimental design. Parental individuals were wild-caught on Jekyll Island, Georgia (GA) and Patchogue, New York (NY). Fertilized eggs for each cross were reared under two different temperatures until larvae reached about 30mm of length. Hatching times and sampling times are given in Table S1. **B)** Temperature-dependent genetic variance in size-at-age (proxy for growth) between individuals used for RNA-seq. The genetic variance of size-at-age was additive at 20°C, with HYB showing intermediate growth. However, size-at-age showed dominance (NY) and/or the influence of maternal effects at 26°C. ANOVA results are highlighted by asterisks: ***: p < 0.05; n.s.: p > 0.05. **C)** Principal component plots for PC1 vs PC2 and PC1 vs PC3 based on all expressed genes. Colours correspond to the crossing scheme in panel A, and temperatures are shown in open (26°C) or closed (20°C) circles. The ellipses give the 95% confidence interval of the data distribution for each cross for both temperatures combined. See Fig. S2 for PC4. **D**) Venn diagram showing the number of unique and shared differentially expressed (DE) genes between NY and GA at 20°C and 26°C. See Fig. S3 for detailed MA-plots.

## Results

### Substantial divergence in gene expression and thermal plasticity across populations

To investigate the gene regulatory mechanisms underlying local adaptation in the Atlantic silverside, we raised F1s from two populations that are locally adapted to different temperature conditions along the North American Atlantic Coast (Hice et al. 2012; Wilder et al. 2020), one low latitude (Jekyll Island, Georgia [GA]) and one mid-latitude (Patchogue, New York [NY]) population and their F1 hybrids (NY x GA; HYB) under two temperatures (20°C and 26°C; Fig. 1a). As expected from prior studies (Conover and Present 1990; Hice et al. 2012), northern NY individuals grew faster than southern GA individuals (Conover and Present 1990; Hice et al. 2012), and growth rates were consistently higher at 26°C than at 20°C (Fig. 1b, Fig. S1). Hybrid individuals showed clear evidence for additive variation in growth with intermediate values at 20°C. At 26°C, the average growth rate of the hybrid group was also in between NY and GA, but not significantly different from the NY group, suggesting maternal effects and/or NY ancestry dominance (see details in Supplementary results; Fig. 1b).

To examine underlying gene regulatory differences, we sequenced whole-body transcriptomes of 42 individuals (6-8 from each population x temperature treatment, Fig 1a) and aligned reads to a linkage-map anchored version of the Atlantic silverside genome (Tigano et al. 2021; Akopyan et al. 2022). After filtering, we retained sufficient information for 18,517 annotated genes. Principal components analysis (PCA) of genome-wide expression patterns revealed a clear separation between individuals from GA and NY along PC1 (PVE_PC1_=24%; Fig. 1c). Most hybrid individuals fell in between the two parental populations along this axis, but overlapped substantially more with the maternal NY population, again suggesting either maternal effects on expression or NY ancestry dominance. In total, more than a quarter of the genes were differentially expressed (DE) GA and NY when compared within a temperature regime, with greater expression divergence at 26°C (5,649 DE genes or 30.6% of all expressed genes) compared to at 20°C (4,655 DE genes or 25.2% of all expressed genes; Fig. 1d, Table S2).

Within each population, temperature also caused substantial differences in gene expression, but, interestingly, along different PC axes, with PC2 (PVE_PC2_=13%) almost completely separating individuals from different temperature-treatments in GA while it is PC3 (11%) and PC4 (8%) that separate temperature-treatments in NY and HYB (Fig. 1d, Fig. S2), indicating that it was a different subset of genes that responded to temperature in the different populations. Temperature-dependence in differential expression patterns between populations for a large number of genes (Fig. S3) indicate the presence of strong genotype x environment interactions (GxE) (Fig. S4, Table S2). Under local adaptation to distinct thermal environments and GxE, we might expect that gene expression responses to temperature differ between populations (Grishkevich and Yanai 2013) (Fig. 2a). Thermal responses did indeed differ between populations, with more DE genes between temperatures in GA (n_GA_ = 2371; 12.8%) than in NY (n_NY_ = 1504; 8.1%) (Fig. 2b, c; Table S2), suggesting stronger thermal responses in GA. The majority of genes showed temperature-specific responses only in one of the populations (n_GA-only_ = 1736, 9.4%; n_NY-only_ = 1235, 6.7%) and a few genes changed expression in opposite directions (n_opposite_ = 38, 0.2%) (Fig. 2b-d, Table S3). Only 269 (1.5%) out of 3278 temperature-responsive genes changed expression in the same direction in GA and NY (Fig. 2b-d, Table S3). Together, these results highlight local adaptation in growth and thermal response in Atlantic silversides.

**Figure 2.**
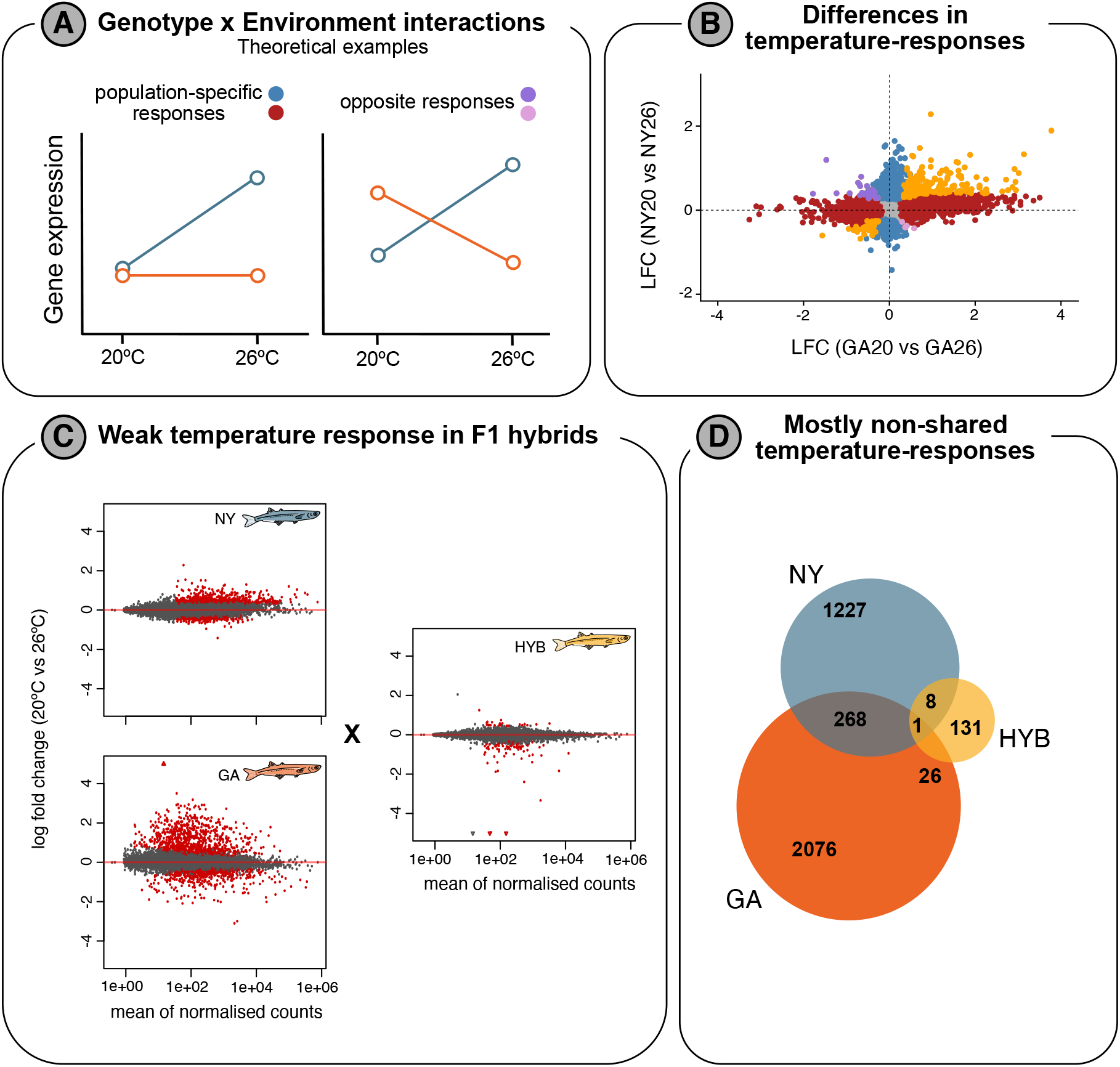
Genotype x environment interactions. **A)** Theoretical examples for genotype x environment interactions (GxE) for individual genes based on temperature-dependent expression profiles within populations. GxE can either include population-specific responses or opposite temperature responses between populations. **B)** Comparison of gene expression differences (log fold change; LFC) between temperature regimes (20°C vs 26°C) in GA and NY. Genes with evidence for GxE are coloured based on the colour scheme in Panel A. Genes with concordant temperature responses between populations, highlighted in orange, do not show any evidence for GxE. **C)** MA-plots highlight that temperature-responses in pure-bred parental F1 populations are stronger in both magnitude and number compared to their F1 hybrids. **D)** Venn diagram showing the large difference and weak overlap in the number of genes showing differential expression across temperatures in the different groups.

### Reduced thermal plasticity in hybrids coupled with temperature-dependent misexpression

Differences in gene expression between hybrids and their parental populations (expression inheritance; Fig. 3a and S5) can reveal gene regulatory incompatibilities between populations, which can manifest in disrupted phenotypes or on the molecular level as gene misexpression in hybrids (transgressive expression) (Coolon et al. 2014; Mack et al. 2016; Mack and Nachman 2017).

**Figure 3.**
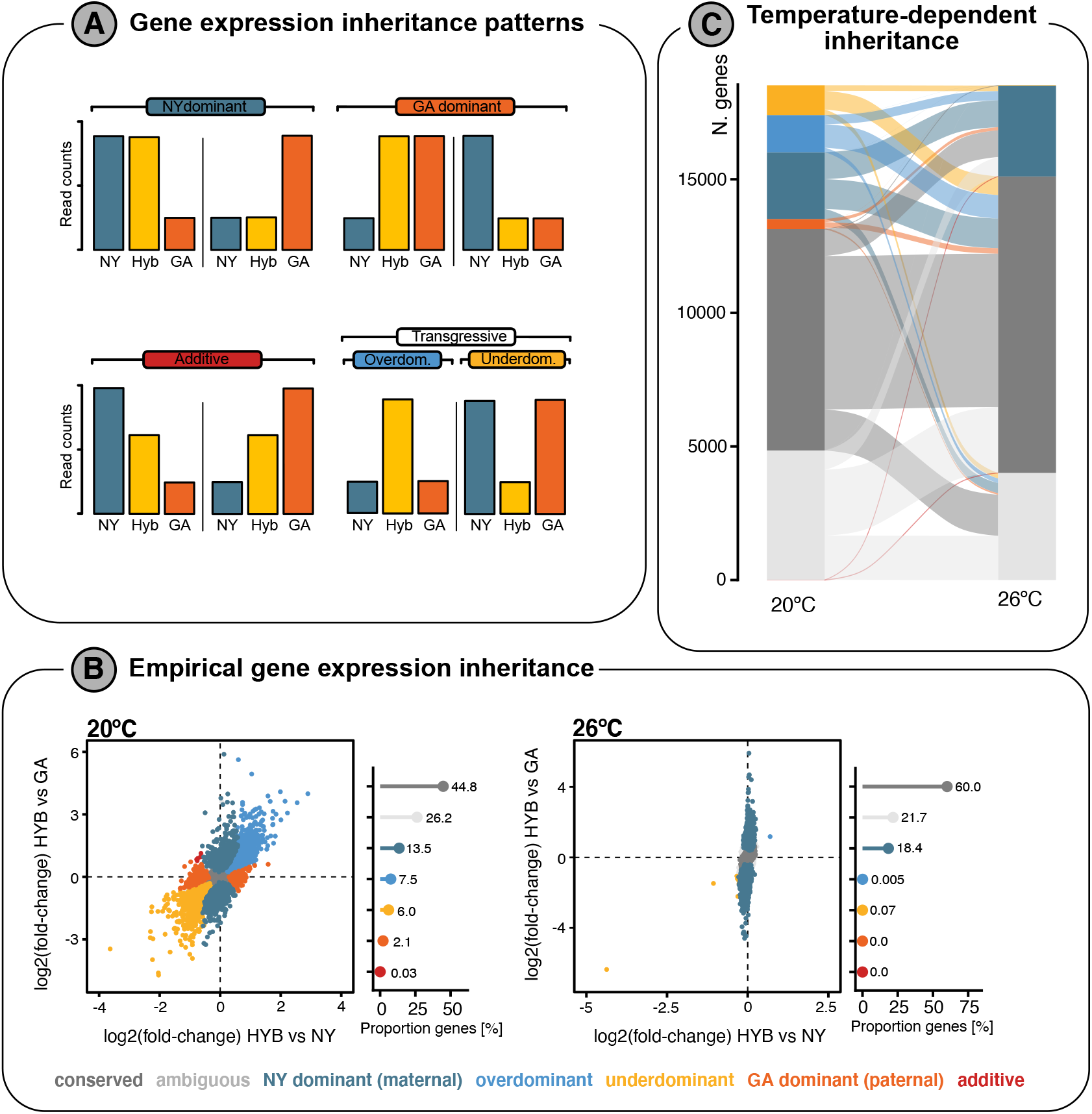
Gene expression inheritance is dependent on temperature. **A)** Cartoon of expected expression patterns among populations under different types of gene expression inheritance (see also Fig. S5). **B)** Comparison of gene expression differences between HYB and both parental populations (NY and GA) to identify inheritance modes for individual genes. The proportion of genes with specific inheritance modes is shown in lollipop plots on the right, with percentages next to each class (see Table S4 for details). Points in all plots are coloured based on the inferred inheritance mode. **C)** Alluvial plot highlighting changes in the number of genes for each inheritance mode between temperature regimes. This plot only includes genes expressed in both temperature regimes.

First, we investigated how thermal responses and gene expression differ between parental and F1 hybrid (HYB) individuals. In stark contrast to both parental populations, a much smaller number of genes (n=166, 0.9%) showed expression differences between temperatures in the HYB group (Fig. 2c, d; Table S3). A large number of genes differed in expression between HYB and GA in a temperature-dependent fashion (20°C = 7068 [38.2%]; 26°C = 3740 [20.2%]). However, we found fewer DE genes between HYB and NY at 20°C (n=3472 [18.8%]) and notably, only 24 DE genes (0.13%) at 26°C, in line with temperature-dependent maternal effects and/or NY ancestry dominance (Fig. 1b, c).

Second, we investigated gene expression inheritance patterns for signs of gene miseexpression in hybrids. Genes that show higher (overdominant) or lower (underdominant) expression in hybrids compared to both parental populations (Fig. 3a), potentially harbour (epi)genetic regulatory incompatibilities between populations (McManus et al. 2010; Groszmann et al. 2013; He et al. 2013; Coolon et al. 2014; Mack and Nachman 2017; Mugal et al. 2020). As expected for recent evolutionary divergences, most genes showed conserved inheritance in hybrids (i.e. no significant difference in expression between hybrids and both parental groups) at both temperatures (60.0% of all expressed genes at 26°C and 44.8% at 20°C; Fig. 3a, Table S4). A relatively large proportion of genes also showed NY (maternal) dominant expression patterns (18.4% at 26°C, 13.5% at 20°C), while we only detected weak GA (paternal) dominant expression for 2.1% of genes at 20°C and none at 26°C, consistent with substantial maternal effects. However, due to the lack of reciprocal crosses we cannot fully distinguish ancestry-dominance from maternal effects. Most strikingly, however, we observed substantial hybrid gene misexpression at 20°C, with over 13.5% of all expressed genes being transgressively expressed in hybrids at 20°C. In contrast, only 0.07% of genes were misexpressed at 26°C (Fig. 3b), with most misexpressed genes at 20°C showing conserved expression patterns at 26°C (Fig. 3b).

To determine the selective forces underlying gene expression evolution between locally adapted populations, we compared DE genes between parental populations and misexpressed genes (McGirr and Martin 2020a). We found that the majority of the misexpressed genes were not differentially expressed between parental populations (20°C: n=1934 [77.2% of all misexpressed genes]; 26°C: n = 8 [61.5%]), in line with stabilising selection maintaining optimal expression levels in the parental populations despite divergence in gene regulatory mechanisms as suggested by the misexpression of those genes in hybrids (Wray et al. 2003). However, a substantial portion of the misexpressed genes were also differentially expressed between NY and GA, particularly at 20°C (20°C: n=571 [22.8% of all misexpressed genes]; 26°C: n = 5 [38.5%]), indicating that expression levels of these genes are potentially under divergent selection between parental populations (Pavey et al. 2010; McGirr and Martin 2020a), although other processes such as drift could also play a role. Furthermore, of the 3009 genes with evidence for GxE interactions (population-specific temperature responses) in at least one of the three groups (Fig. 2b), 677 (24.5%) were misexpressed in hybrids in at least one temperature regime (670 at 20°C and 8 at 26°C, with one shared), which is more than expected by chance (HGT_20_°_C_: p = 3.17e-31; HGT_26_°_C_: p = 0.00065). Together, these results suggest that gene regulatory mechanisms have rapidly diverged in a temperature-dependent manner between populations despite ongoing gene flow.

### Temperature-dependent gene regulatory architectures

To better understand whether the expression divergence across populations is primarily driven by changes in *cis*-regulatory (e.g. enhancers) or *trans*-regulatory elements (e.g. transcription factors), we mapped differences in regulatory mechanisms between GA and NY using allele-specific expression (ASE) analyses in hybrids at each temperature. F1 hybrids carry a GA (southern) and NY (northern) copy of each chromosome within a shared *trans* environment (the hybrid cell), hence, any ASE bias at a particular gene can be attributed to the divergence in *cis*-regulatory elements, rather than *trans*-acting factors (Wittkopp et al. 2004; Coolon et al. 2014). If no ASE bias is present in hybrids but parental populations show divergent expression, *trans*-acting factors likely cause the regulatory divergence (Wittkopp et al. 2004; Coolon et al. 2014).

Overall, of the 820 genes with allele counts available for ASE analyses, we found evidence for ASE at 134 genes (FDR < 5%) and 58 genes (FDR < 5%) at 20°C and 26°C, respectively (Fig. 4, Fig. S6). Expression differences between GA and NY were more often associated with *all-trans* differences compared to *cis*-regulatory divergence (*all-cis*) at both temperatures (Fig. 4a), which is generally expected for rapid intraspecific divergences (McGirr and Martin 2020b). Of all expressed genes with heterozygous SNPs, 8.4% (52 out of 619 genes) and 15.8% (102 out of 644 genes) were identified as *all-trans* regulated at 20°C and 26°C, respectively. In comparison, *all-cis* regulatory divergence at both temperatures was less prevalent, with 21 genes (3.4%) at 20°C and 16 genes (2.5%) at 26°C (Fig. 4, Fig. S6). This was significantly lower than the proportion of *all-trans* regulated genes (26°C: *X*^2^=80.91, df=1, p<2.2e-16; 20°C: *X*^2^=14.1, df=1, p=8.629e-05).

**Figure 4.**
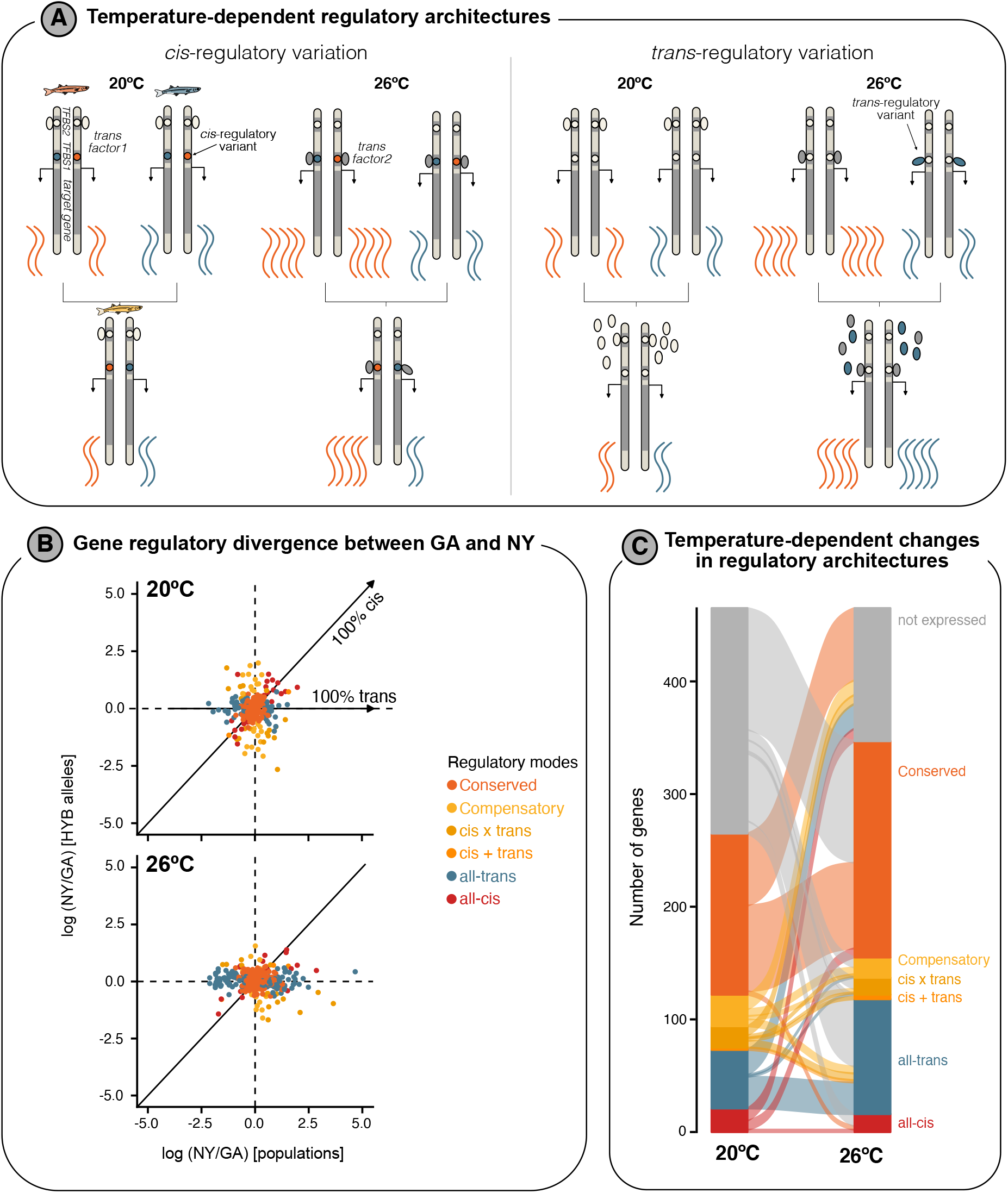
Temperature-dependent gene regulatory architectures. **A)** Illustrations highlighting how regulatory architectures can diverge in *cis* and *trans*, and how this might affect expression levels in hybrids, and importantly how environmental differences might lead to different inferred regulatory modes, e.g. conserved at 20°C but *cis*- or *trans*-regulatory divergence at 26°C. TFBS = Transcription factor binding site. **B)** Scatterplots show differences in expression between parental populations versus expression differences between maternal and paternal alleles in hybrids at 20°C (top) and 26°C (bottom). Dots are coloured by their inferred regulatory mode and represent individual genes. Expected expression patterns for all-cis and all-trans regulated genes are highlighted by solid lines/arrows. **C)** Changes in the number of genes for each inferred regulatory mode between temperature regimes are illustrated in this alluvial plot. Genes that are not expressed under one temperature regime are shown in grey.

In contrast to theoretical expectations of higher context-dependencies of *trans*-regulatory factors (Cutter and Bundus 2020), we found that the proportion of genes consistently *all-cis* regulated across temperatures (6.1% of *all-cis* regulated genes were *all-cis* regulated at both 20°C and 26°C) was lower compared to the proportion of consistently *all-trans* regulated genes (24.8% of *all-trans* regulated genes were *all-trans* regulated at both 20°C and 26°C). However, nine genes that were *all-trans* regulated at one temperature showed evidence for an interaction with *cis*-regulatory evolution (‘Compensatory or reinforcing regulatory architectures’, see Supplementary text) at the other temperature (Fig. 4c).

We further investigated if potential regulatory incompatibilities between co-evolved *trans*- and *cis*-factors are associated with misexpression in hybrids (McGirr and Martin 2020a). Of the 85 and 90 misexpressed genes that were available for our ASE analysis at 20°C and 26°C, respectively, only seven showed evidence for compensatory evolution at one temperature (7.8 - 8.2% of genes), supporting the action of stabilising selection acting on optimal expression levels of these seven genes in the parental populations.

### Disruption of temperature-sensitive gene expression networks in hybrids

In general, genes do not act independently but rather interact within networks of co-expressed genes. The properties of co-expression networks can also evolve and low preservation of networks and/or their disruption in hybrids might point toward regulatory changes involved in local adaptation (Filteau et al. 2013). We used WGCNA (Langfelder and Horvath 2008) coupled with module preservation analysis (Langfelder et al. 2011) to infer and compare co-expression networks between the NY and GA populations and their hybrids. Module preservation analysis was used to test whether gene composition and among-gene connectivity properties of modules in each pure population network were maintained in the hybrid network. We expected that pure population modules that are weakly preserved in the hybrid network represent potential gene regulatory pathways that are misexpressed.

WGCNA clustered 18,526 genes into roughly equal numbers of modules within NY, GA, and hybrid networks (NY: 23 modules; GA: 28 modules; hybrid: 22 modules; Table S6-7). While most modules were at least moderately preserved between NY and GA (Z_*summary*_ > 10), we detected 7 (30.4%) and 13 (46.4%) modules in NY and GA, respectively, that were not strongly preserved in the other population (Table S6-7). We used analysis of covariance (ANCOVA using body length as a covariate) to determine the effect of rearing temperature on module eigengene (module expression) and separately, Pearson correlations to test for a relationship between eigengene and size-at-age (as a proxy for growth). We found that 5 and 11 modules in the GA (Table S7) and NY (Table S6) networks, respectively, showed an effect of temperature, and 3 and 5 modules were significantly associated with size-at-age in GA and NY, respectively (Fig. 5a). Three temperature-sensitive modules in NY, including one growth-associated module (brown NY module), were only weakly preserved in GA (Table S7). Similarly, two temperature-sensitive modules in GA were only weakly preserved in NY, one of which (tan module) was also significantly correlated with size-at-age (Table S6).

**Figure 5.**
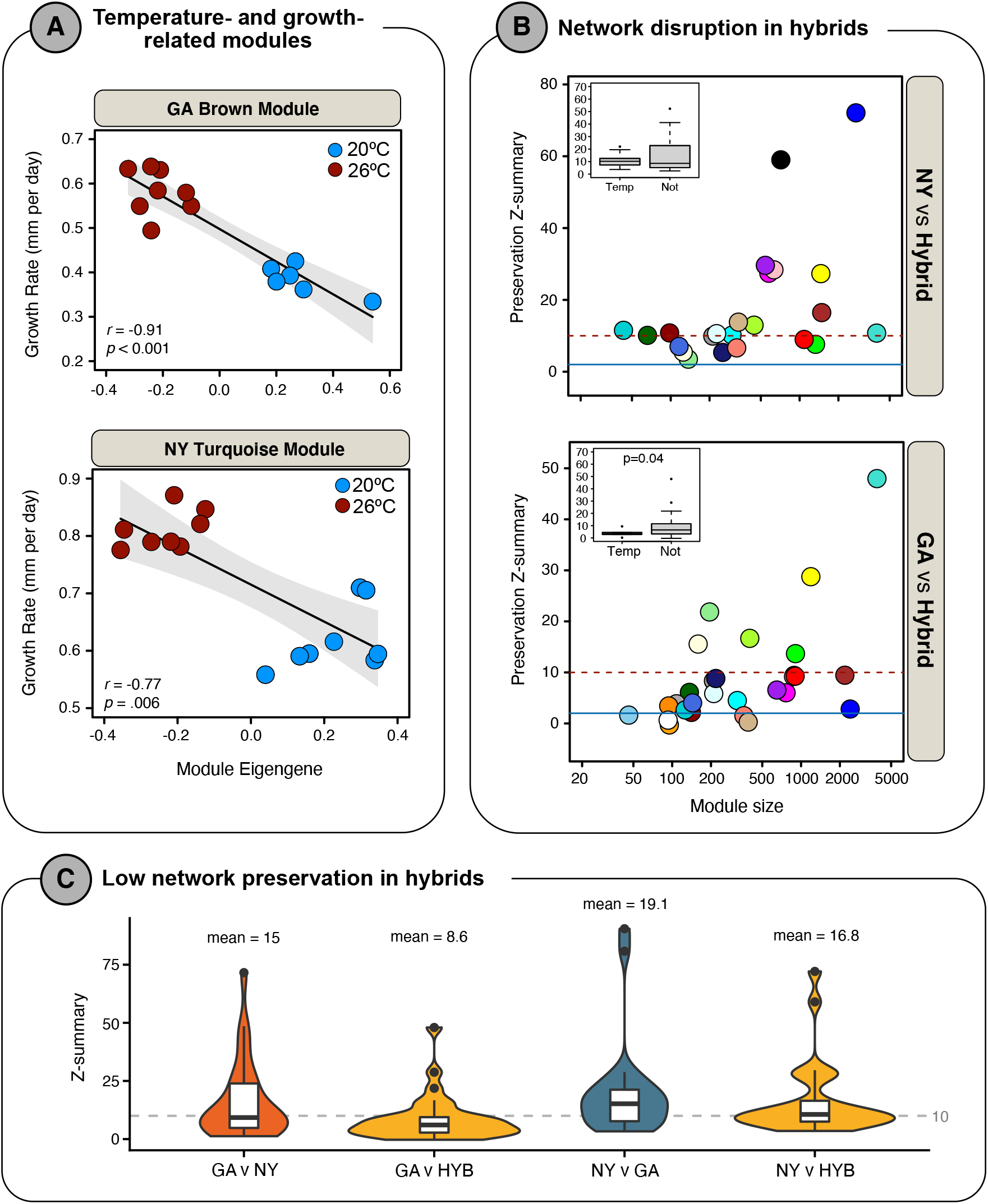
Disruption of gene expression networks. **A)** Correlation between growth rate between module eigengene expression for two temperature-responsive modules, the Brown module in GA and the Turquoise module in NY. **B)** Module preservation composite Z_summary_ scores for each of two comparisons between NY and GA networks and the hybrids. Each colored point represents a module identified in the pure GA network (A) or pure NY network (B). Z_summary_ scores are plotted against the size of each module in numbers of genes (see Tables S6 and S7). Insets are the boxplots of Z_summary_ scores in each comparison for modules responsive to temperature (Temp.) or not responsive to temperature, as identified in ANCOVA (Tables S6 and S7). **C)** Network preservation, given as Z-summary score, for each co-expression network comparison shown as boxplots, with the distribution of values shown as violin plots. HYB networks are generally less well preserved than pure networks. The grey, dashed line shows a Z-summary score of 10, below which modules are considered weakly preserved. Mean values for each comparison are also given.

To determine which biological processes are potentially divergent between these populations, we performed gene ontology enrichment analyses on temperature-sensitive and growth-associated modules. However, functional enrichment analysis of temperature- and growth-associated modules was difficult to interpret, likely because we analyzed expression in whole animals, capturing profiles across many different cell and tissue types. We did, however, find that growth- and temperature-associated modules were both enriched for metabolism of fat as fuel (Supplementary file S1). Expression of these modules was negatively correlated with size-at-age and temperature (Fig 5a), suggesting that fatty acid metabolism is downregulated when fish are reared in warm environments (where size-at-age is highest). The temperature- and growth-related modules that were only weakly preserved between NY and GA (brown NY module, tan GA module), were additionally also functionally enriched for a wide range of processes associated with growth, development and cellular differentiation (Supplementary file 1), processes that are known to differ between those populations (Yamahira and Conover 2002; Hice et al. 2012). Furthermore, the brown NY module was also enriched for processes related to thermal response (Supplementary file 1).

We next tested whether pure population modules were disrupted in hybrids, which would support the divergence and incompatibility of the underlying gene regulatory mechanisms between NY and GA observed on the single gene level (Fig. 4). Indeed, the majority of pure population modules were not well preserved in the hybrid network (Fig. 5b, c). This pattern was most pronounced in GA where 79% of modules were weakly (Z_*summary*_ < 10) or not-at-all (Z_*summary*_ < 2) preserved in hybrids (Fig. 5B; Table S7). Modules in the NY-hybrid comparison were better preserved than in the GA-hybrid comparison, although preservation was still generally low, with 57% of modules being weakly or not-at-all preserved (Fig. 5; Table S6). Temperature-responsive co-expression modules were generally less preserved than modules not influenced by temperature, though this result was only significant in the GA vs hybrid comparison (Fig. 5b, see inserts). Of the growth-associated modules, only two (out of five) from the NY network were well-preserved in hybrids (Table S6), and none from the GA network were (Tables S7). Moreover, pure population modules tended to be less preserved in hybrids than each pure population network in comparison to each other. For example, over 2X as many GA modules were preserved in the NY network than in the hybrid network (48% vs. 21%; Table S6), with the mean and median Z_*summary*_ scores being consistently higher in pure population comparisons compared to comparisons involving hybrids (Fig 5c). This observation might be explained by the compensatory evolution of underlying gene regulatory mechanisms, meaning that co-expression patterns are under stabilising selection within each lineage despite the divergence of the underlying regulatory mechanisms. In line with the presence of gene misexpression (Fig. 4) and divergence in regulatory mechanisms (Fig. 5), which we also detected for central hub genes in co-expression modules (Supplementary results), these results support the presence of stark regulatory divergence between locally adapted silverside populations despite gene flow.

### Temperature-dependent contribution of inversions to regulatory divergence

To begin exploring how the large inversions that show highly elevated differentiation between populations may affect gene regulatory patterns, we examined whether divergently regulated genes were enriched within these inversions. If inversions play a predominant role in gene regulatory divergence, we expect enrichment for DE genes and misexpressed genes within inversions segregating in our dataset on chromosome 4 (*inv4*), chr.7 (*inv7*), chr.8 (*inv8*), chr.18 (*inv18*), chr.19 (*inv19*) and chr.24 (*inv24*) (Fig. S7-8). Overall, 527 (11.3%) and 661 (11.7%) of all DE genes, and 248 (10.2%) and 5 (41.6%) of all misexpressed genes, were located inside segregating inversions at 20°C and 26°C, respectively. These relatively small proportions suggest that inversions are not the predominant source of regulatory variation, at least not directly. However, the inversions only cover a relatively small proportion of the genome (∼16%) (Akopyan et al. 2022), so we tested if the proportion of DE and misexpressed genes was higher within individual inversions compared to the collinear genome. Indeed, DE genes were significantly enriched within *inv4* and *inv18* at 20°C, but only within *inv18* at 26°C (Table 1). In contrast, misexpressed genes were enriched within all inversions at 20°C, and within *inv4, inv8*, and *inv18* at 26°C (Table 1).

**Table 1.**
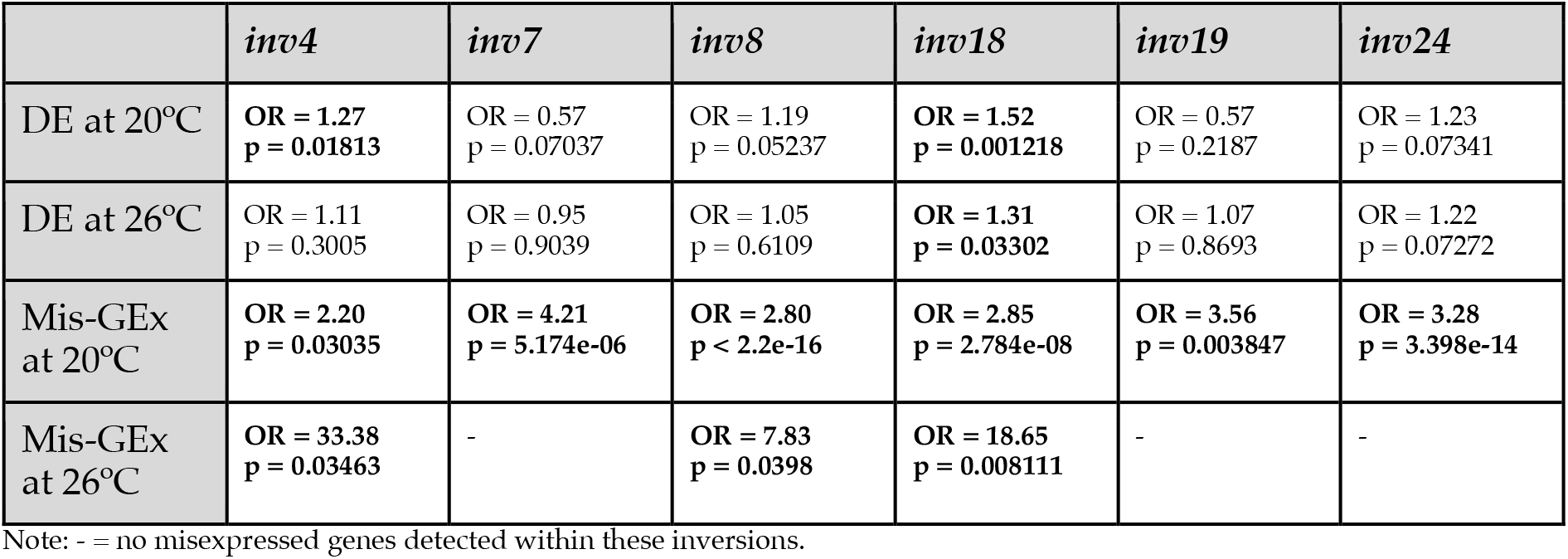
Inversion enrichment results. Shown are odds ratio (OR) and p-values from Fisher’s exact tests, comparing the proportion of DE genes or misexpressed genes within inversions compared to the proportion of DE or misexpressed genes in the collinear genome.

Furthermore, *inv18* and *inv24*, the two inversions that show the strongest differentiation in allele frequencies between our populations (Akopyan et al. 2022), were also enriched for genes belonging to certain GA and NY gene modules. Genes belonging to the “tan” NY-module and “darkorange” GA-module were enriched inside *inv18*, and genes from the “darkgreen” NY-module and the growth-related “brown” GA-module (Fig. 5a) inside *inv24* (Table S7). All of these modules showed temperature-dependent expression, and three of the four modules were only weakly preserved in the GA-NY and/or pure-hybrid comparison (Table S6-7), suggesting that the enrichment within the most strongly divergent inversions on chromosomes 18 and 24 is potentially associated with the disruption of these networks, perhaps in a temperature-dependent manner.

## Discussion

Despite our growing understanding of the genomic changes underlying local adaptation with gene flow, which are often associated with chromosomal inversions, the underlying molecular mechanisms, and the contribution of putatively locally adaptive inversions to gene regulatory evolution remain largely unknown. Here, we identify the gene regulatory mechanisms underlying local adaptation under pervasive gene flow in a marine fish that harbours multiple locally adaptive inversions, the Atlantic silverside. Overall, differential gene expression and differences in temperature-responses are substantial between locally adapted populations. Misexpression and disruption of co-expression modules is pervasive at colder rearing temperatures, yet less pronounced at warmer temperatures, pointing to a role of condition-dependent regulatory incompatibilities in local adaptation. Misexpressed genes are strongly enriched within chromosomal inversions, suggesting that these structural variants have accumulated incompatible alleles. Contrary to our expectation, the substantial divergence in gene expression is more strongly driven by *trans*-than *cis*-regulatory divergence, although regulatory mechanisms were often temperature-dependent. Overall, we provide evidence for an important role of gene regulatory divergence and regulatory incompatibilities in local adaptation with gene flow in a highly connected marine fish.

### Strong divergent genome-wide expression and temperature responses between locally adapted populations

Gene expression is highly divergent between locally adapted Atlantic silverside populations despite gene flow between them (Wilder et al. 2020), with over 30% of all genes being differentially expressed between populations (Fig. 2a). Such levels of expression divergence are extremely high compared to estimates from other studies focused on intraspecific comparisons, which largely range from 0.45% to a maximum of 20% genes being differentially expressed (Barreto et al. 2015; Juneja et al. 2016; Hanson et al. 2017; Velotta et al. 2017; Mack et al. 2018; Fischer et al. 2021; Jacobs and Elmer 2021). They are more comparable to strong interspecific divergences between *Drosophila spp*. (25-35% of genes; (Coolon et al. 2014)), highlighting strong regulatory differences between Atlantic silverside populations. We note though that in contrast to several of the above-mentioned studies, our analyses measured expression across whole juvenile organisms rather than individual tissues, which could lead to a higher proportion of differentially expressed genes. Tissue-specific analyses in Atlantic silversides will be needed to better understand the molecular mechanisms and extent of divergence across different organs.

Furthermore, the strong divergence in thermal plasticity between populations (largely non-overlapping sets of genes responding to temperature shifts in the different populations) suggests local adaptation in temperature-response between NY and GA on the molecular level (Conover and Present 1990; Hice et al. 2012), which has been observed in other systems occupying divergent thermal habitats (Campbell-Staton et al. 2021). The stronger divergence in gene expression between temperature regimes in GA compared to NY suggests that thermal plasticity is stronger in southern silverside populations inhabiting warm environments with less pronounced seasonal variation. This pattern is opposite to Atlantic cod, where cold-adapted populations seem more sensitive to temperature changes (Hutchings et al. 2007). The stark molecular divergence in temperature-responsive pathways is further supported by the low preservation of many temperature-responsive gene co-expression networks between NY and GA (Fig. 5; Table S6-7). The weak thermal plasticity and low preservation of temperature-responsive co-expression modules in F1 hybrids suggests the disruption of temperature-responsive pathways in hybrids, as observed in other fish (Oomen et al. 2021; Payne et al. 2021). If this disruption of temperature-responsive pathways leads to lower fitness in wild hybrids, it might provide a potential explanation for the maintenance of local adaptation despite ongoing gene flow. However, we have so far only focused on gene expression, but other regulatory processes, such as alternative splicing, might also play important roles in thermal adaptation in silversides, and could compensate for disrupted expression patterns (Healy and Schulte 2019; Verta and Jacobs 2022). In contrast, there is the possibility that hybrids show higher thermal tolerance than parental populations (Pereira et al. 2014) and further experimental studies are needed to determine the physiological effects of hybridization in this system.

### Maintenance of local adaptation through hybrid gene misexpression

Regulatory incompatibilities, genetic or epigenetic, between populations or species can lead to the misexpression of genes in hybrids, even at early stages of divergence (Renaut et al. 2009; Barreto et al. 2015; McGirr and Martin 2020a; Mugal et al. 2020; Moran et al. 2021). Such regulatory incompatibilities can facilitate the maintenance of local adaptation despite gene flow, if the resulting misexpression has negative fitness effects (Filteau et al. 2013; McGirr and Martin 2020a). Atlantic silversides show surprisingly strong misexpression in hybrids (13.5% of all expressed genes) at lower temperatures (20°C) compared to other examples of intraspecific clinal adaptation, such as in the copepod *Tigriopus californicus* that shows hybrid misexpression in 1.2% of genes (Barreto et al. 2015), and even slightly higher than levels of misexpression in crosses of young species pairs, e.g. lake whitefish (10%) (Dion-Côté et al. 2014) or Caribbean pupfish (up to 9.3%) (McGirr and Martin 2020a). While misexpression in hybrids could result from aberrant development (Mack and Nachman 2017), we did not detect any stark developmental differences (e.g. in hatching time) between pure and hybrid crosses reared at the same temperature, supporting the role of regulatory incompatibilities. However, the very low proportion of misexpressed genes at 26°C highlights how misexpression can be highly condition-dependent, and that misexpression is contributing more strongly to local adaptation under more challenging environmental conditions, in this case lower temperatures.

Most misexpressed genes are thought to be under stabilising selection in parental populations (Wray et al. 2003; Moran et al. 2021), resulting in similar expression levels between populations. Yet, we found several hundred misexpressed genes that were differentially expressed between GA and NY populations in Atlantic silversides, suggesting that their expression levels are under divergent selection and putatively involved in local adaptation (Pavey et al. 2010; Kulmuni and Westram 2017; McGirr and Martin 2020a). These genes are potentially involved in the divergence of locally adaptive phenotypes. Future work should focus on determining the molecular mechanisms of hybrid gene misexpression on the tissue level and determine the phenotypic and fitness effects of misexpression to better understand their contribution to local adaptation.

Furthermore, the low preservation of co-expression modules, including temperature-responsive and growth-correlated modules (Fig. 5, Table S6-7), could partially be explained by potential genetic incompatibilities and the resulting misexpression of central regulatory genes within modules (Supplementary text). The disruption of growth-related modules in hybrids could be related to the observed difference in growth rate in hybrids compared to the parental populations, particularly intermediate growth rates at 20°C. This difference in growth rate has potentially negative fitness effects in their parental habitats, particular in northern habitats, as growth rates show countergradient variation, with selection for higher intrinsic growth rates in northern populations to compensate for shorter growing seasons (need to reach large size before winter to survive), and potential selection on lower growth rates in the south as fast growth can have potentially negative effects on swimming abilities and predator escape (Billerbeck et al. 2001; Lankford et al. 2001). Thus, while intermediate growth rates may be adaptive at central locations of the thermal cline, it might be maladaptive in either parental habitat.

### Inversions and local adaptation under gene flow

In species with largely concentrated genomic architectures, e.g. due to inversions, one might expect that differentially regulated genes are located within inversions due to the accumulation and maintenance of locally adaptive genetic variation through reduced recombination (Stevison et al. 2011; Mérot et al. 2020). In a few systems, researchers indeed found that inversions indeed are enriched for DE genes (Marquès-Bonet et al. 2004; Cassone et al. 2011; Fuller et al. 2016; Berdan et al. 2021), suggesting that inversion-linked *cis*-regulatory variation is the main cause for divergent gene expression (Crow et al. 2020; Berdan et al. 2021). However, most DE genes were located outside inversions in Atlantic silversides. This contrasts with patterns of genetic differentiation observed between NY and GA that show most pronounced differentiation within inversions, particularly *inv18* and *inv24* (Fig. S7; Akopyan et al. 2022). Thus, inversions do not only have *cis*-effects on expression but might have more widespread genome-wide effects on expression through *trans*-specific effects, e.g. through the differential expression of transcription factors linked to inversions (Naseeb et al. 2016; Said et al. 2018). However, inversions also show evidence for *cis*-effects on expression, as supported by the enrichment for DE genes (*inv4, inv18*) and genes from growth- and temperature-associated co-expression modules (*inv18* and *inv24*). In particular, *inv24* has been previously associated with body size evolution in Atlantic silverside (Therkildsen et al. 2019). Together, this suggests that *inv18* and *inv24* harbour adaptive *cis-*regulatory genetic variation underlying locally adaptive traits, but that inversions in general have strong inversion-linked *trans*-regulatory effects on genome-wide gene expression.

In addition to driving differential expression, inversions can potentially contribute to population divergence by reducing hybrid fitness through the accumulation of genetic incompatibilities that lead to hybrid gene misexpression and the disruption of regulatory networks (Navarro and Barton 2003; Feder et al. 2014; Mugal et al. 2020). In contrast to DE genes, all inversions are enriched for misexpressed genes in Atlantic silversides, especially at 20°C. Thus, inversions likely contribute to the maintenance of local adaptation under gene flow via accumulation of epigenetic or genetic incompatibilities that lead to aberrant expression in hybrids, and potentially reducing their fitness in their parental habitats.

### Regulatory mechanisms are highly temperature-dependent

In contrast to protein-coding changes, gene regulatory variation offers the possibility to minimise fitness trade-off across environments by adjusting expression levels between divergent habitats, leading to potential conditional neutrality (Bono et al. 2017; Wadgymar et al. 2017; Gould et al. 2018). While we cannot determine if differences in the observed gene expression divergence between temperatures leads to differences in fitness impacts across environments, we found evidence for GxE for thousands of genes in the Atlantic silverside. Furthermore, the drastic difference in the number of misexpressed genes in hybrids between environments, with nearly no misexpression at 26°C but pervasive misexpression at 20°C, suggests that genetic incompatibilities and fitness effects might be largely invisible at 26°C. This fits observed growth patterns in hybrids, which are similar to the maternal NY population at 26°C (Fig. 1b), suggesting no strong negative effects on growth at higher temperatures, compared to the intermediate growth rate at 20°C. Since we only used unidirectional crosses (NY x GA) in this experiment, due to high mortality of our GA x NY cross, we cannot disentangle maternal from other effects. In general, colder temperatures are thought to be more challenging for Atlantic silversides, and genetic incompatibilities and the resulting fitness effects might be more prevalent in suboptimal habitats (Bomblies et al. 2007). This asymmetry in misexpression might shape the outcome of hybridization between populations, e.g. with potentially less severe negative fitness impacts at warmer temperatures.

Contrary to theoretical expectations (Cutter and Bundus 2020), we found that *cis*-regulated genes were more often temperature-dependent than *trans*-regulated genes, suggesting that *cis*-regulatory elements, e.g. transcription-factor binding-sites, are potentially more temperature-dependent than the expression of transcription factors in Atlantic silversides. This inference might be biased by the small number of genes available for ASE analysis, although it is supported by context-dependent *cis*-regulatory architectures in other species (Gould et al. 2018; York et al. 2018; Cutter and Bundus 2020; Findley et al. 2021). Overall, this highlights the need to study gene regulatory mechanisms across relevant contexts, e.g. developmental stages or environments, and genotypes/divergent populations, to better understand the gene regulatory mechanisms associated with local adaptation across conditions.

### Limitations and Conclusions

Together, this work highlights that regulatory mechanisms and thermal responses can rapidly diverge between locally adapted populations in the face of gene flow. This can lead to widespread misexpression and the disruption of gene co-expression networks in hybrids under certain environmental conditions, suggesting the presence of regulatory incompatibilities, with a strong role of inversions in maintaining such incompatibilities. However, this work is only based on unidirectional crosses from two populations, a single developmental stage and whole-body expression patterns, limiting the inference power. Despite these limitations, this study provides an important starting point for better understanding the role of gene regulatory divergence in local adaptation with gene flow, and for determining how inversions shape the molecular mechanisms underlying local adaptation.

## Methods

### Common garden and plasticity experiment

Wild adults were caught at spawning time using beach seine nets at Jekyll Island, Georgia (GA; 31°03’N, 81°26’W) and Patchogue, New York (NY; 40°45’N, 73°00’W) in spring 2017. Individuals were transported live to the Rankin Seawater Facility at the University of Connecticut’s Avery Point campus. A full reciprocal crossing design was set up by strip-spawning males and females in batches onto mesh screens submerged in plastic dishes in seawater. We created all four possible F1 crosses: NY♀ x NY♂ (NY), NY♀ x GA♂ (NYxGA), GA♀ x NY♂ (GAxNY), and GA♀ x GA♂ (GA), with each cross created from a mix of gametes from ∼40 females and ∼40 males from the specified population. Fertilised eggs were kept in 20L rearing containers placed in large temperature-controlled water baths at constant salinity (30 psu) and photoperiod (15L:9D). We split the fertilized eggs of each pure cross (NY and GA) into four batches, and hatched and reared two batches per cross at 20°C and two batches at 26°C. The hybrid crosses were each split into two batches, with one batch for each crossing direction incubated at either 20°C or 26°C. The two temperatures, 20°C and 26°C, were chosen to reflect the common rearing temperatures in the wild at each of the parental spawning locations (NY and GA), respectively. Individuals were reared to an approximate total length of 30 mm (Fig. S1), with the rearing durations (from 37 - 87 dpf; Table S1) differing between populations and temperature regimes because of strong variation in growth rates between treatment groups (Fig. 1a) (Yamahira and Conover 2002; Hice et al. 2012). We sampled individuals at the same size rather than the same age (days since fertilisation) to allow comparison of individuals that had reached a similar developmental stage across all treatments. In the GAxNY (GA mother) cross, most individuals died at 26°C, and all died at 20°C and thus we could not include this cross in the present study. From each of the remaining crosses, we randomly selected 6-8 individuals for RNA-sequencing.

We approximated individual growth rates for each sequenced individual by dividing the total length (in mm) at sampling by their age (in days from hatching to sampling). To compare growth rates across populations and temperatures, we performed an analysis of variance (*ANOVA*) using the *aov* command in R, including population, temperature and their interaction as terms. Furthermore, we performed a Tukey Honest Significant Differences test on the ANOVA results (*TukeyHSD* function in R) to post-hoc estimate p-values between individual groups [population * treatment].

### RNA-seq & raw data processing

Total RNA was extracted from whole larvae (n=42; Table S1) using the ZymoResearch Direct-zol Miniprep RNA plus kit, following homogenization in Trizol using a pestle. An in-column DNAse I treatment step was performed. RNA quantity was determined using the HS Assay kit for the Qubit 3.0 fluorometer (Life Technologies, Carlsbad, CA) and quality was assessed using a Fragment Analyzer (Agilent, Santa Clara, CA) at the Cornell University Biotechnology Resource Centre. RIN values ranged from 5.3 to 8.3, with an average RIN of 6.9. RNA-seq libraries were prepared at BGI Genomics using the stranded Illumina TruSeq mRNA sequencing kit with Poly-A selection and each library was sequenced to an average of 37.1M 150bp paired-end reads (± 0.194M s.d.) using an Illumina HiSeq 4000 sequencer at BGI.

Raw sequencing data were aligned to the linkage-map anchored *Menidia menidia* reference genome (Tigano et al. 2021; Akopyan et al. 2022) using *STAR* (Dobin and Gingeras 2015) with default settings. The chromosome-scale genome used for this analysis was improved by anchoring the Atlantic silverside reference genome v1 (Tigano et al. 2021) to a female GA linkage map (Akopyan et al. 2022) (see Supplementary methods for details; Table S8). Duplicates were marked using the picard tools and read count tables for each annotated coding sequence (CDS) and each individual were generated using *HTSeq-count* for reads with a minimal alignment quality of 20 (*-a 20*).

### Gene expression analyses

Differential gene expression analyses were performed using the R-package *DESeq2* (Love et al. 2014). For the initial analyses and visualisation, we analysed all samples together, retaining all genes with a minimum of 42 counts per million (∼1 cpm per individual) and using the following model: ∼ population + temperature + population * temperature + body length; with populations being NY, GA or HYB (NYxGA), temperature either 20°C or 26°C, and length the total body length of each individual at the time for sampling. Principal components analyses were performed using the *pcaMethods* R-package (scaling = “none”, centre = TRUE) on *rlog*-transformed read counts (Stacklies et al. 2007). We identified significantly differentially expressed genes between populations across temperatures using *DESeq2* with an alpha threshold (false-discovery rate) of 0.05 and used the ‘ashr’ shrinkage methods for ranking genes by effect size (log fold change). We also tested for the effect of body length on patterns of gene expression across all samples to test if gene expression variation was driven by variation in size/developmental stage, but no significant effect of length was detected for any of the genes (p>0.05).

Due to the strong effect of temperature on expression patterns (see PCA, Fig. 1c, Fig. S2), we identified differentially expressed genes separately for each rearing temperature using *DESeq2* with the following model: ∼ population + body length. We only included genes with a minimum count of 20 and 22 counts per million across all individuals at 20°C and 26°C, respectively.

To determine genotype x environment interactions (GxE), we identified the sets of genes that showed differential expression between temperatures in each of the two pure populations (GA vs NY), defining genes with evidence of GxE as those falling into one of the following two categories (Fig. 2a): 1) only one population, i.e. genotype, shows significant differential expression between temperatures (FDR < 0.05) (population-specific response), or 2) both populations show significant differences between temperatures, but in opposite directions (opposite response). We also identified genes that showed similar (conserved) responses to temperature in both populations.

Lastly, we determined the inheritance mode for each gene based on the difference in gene expression between the pure parental populations and the hybrids following the criteria in Coolon *et al. (Coolon et al. 2014)* (Fig. 3a, Supplementary methods). We combined genes with *overdominant* and *underdominant* expressions as ‘misexpressed’. We performed the statistical comparison of expression for each gene across replicates (individuals from the same group [population * temperature]) using DESeq2 as outlined above.

### Allele-specific expression analysis

To determine differences in regulatory modes between populations, we performed allele-specific expression (ASE) analyses across hybrids. To reduce the impact of allele-specific mapping bias on our analyses, we first produced a reference genome with all fixed and nearly fixed SNPs between populations masked. We did this by first calling SNPs from the genome-aligned RNA-seq data using GATK v.3.8 (McKenna et al. 2010) using the HaplotypeCaller pipeline. We only retained bi-allelic SNPs with a genotype quality above 30, minimum depth of 5x and maximum depth of 52, minor allele frequency above 5% and with less than 25% missing data. We determined allele frequencies in each population across temperatures using vcftools v.0.1.16 and identified fixed and nearly fixed sites based on allele frequency difference (AFD) following (Berner 2019). We then used the *ASEr* pipeline (Combs and Fraser 2018) (https://github.com/TheFraserLab/ASEr) to mask all heterozygous sites with AFD values above 0.95th percentile of the empirical distribution (AFD > 0.392). While interspecies comparisons typically focus on only fixed genetic differences between species (McManus et al. 2010; Coolon et al. 2014), we also included strongly differentiated SNPs in our intraspecies comparison, as fixed sites are relatively rare due to the strong gene flow and differentiated SNPs are also functionally important. Furthermore, our ASE analysis approach simultaneously compares many replicates in a population-level approach (Wang et al. 2020) rather than focusing on individual parent-offspring pairs, thus increasing the usefulness of highly differentiated, but non-fixed, SNPs. Subsequently, we remapped all RNA-seq data for hybrid individuals back to the masked reference genome, keeping only uniquely mapping reads, and produced allele-specific haplotype counts for all heterozygous SNPs using the *phaser* pipeline (Castel et al. 2016). In brief, we phased the VCF file with all masked heterozygous sites using *beagle* v.4.1 with default settings and used the phased VCF and re-mapped hybrid bam files as input for *phaser*, only keeping sites within genes with a minimum base-quality of 10 and reads with a minimum mapping quality of 255, resulting in allele counts for 820 genes. To test for potential mapping bias, we confirmed that the distribution of haplotype counts is centred around zero (Fig. S9). Lastly, to test for allele-specific expression and classify genes by regulatory mode, we followed the statistical approach by Wang et al. (Wang et al. 2020) based on the criteria by Coolon et al. (Coolon et al. 2014) (Supplementary text). Compared to Coolon et al. (Coolon et al. 2014), Wang et al. (Wang et al. 2020) made use of biological replicates, as we have in our study, and analysed allele-specific expression by fitting Negative binomial generalized linear models and Wald statistical tests using *DESeq2*. In brief, we compared NY/GA alleles in hybrids using the transTest with the following formula design: ‘∼0 + Gen * Origin’, with ‘Gen’ denoting the allele (GA or NY) in hybrids and ‘Origin’ denoting whether reads were from the parental individuals or hybrids. After classifying genes with low read counts as ‘uninformative’, we classified the remaining genes as *‘ambiguous’, ‘conserved’, ‘compensatory’, ‘all-cis’, ‘all-trans’*, ‘cis+*trans’* or *‘cis×trans’* (Supplementary text). We performed the ASE analysis independently for each temperature-regime.

### Co-expression network analyses

We used module preservation analysis (Langfelder et al. 2011) to test for evolved, population-specific gene regulatory responses to rearing temperature. WGCNA (Langfelder and Horvath 2008) produced gene co-expression networks for each population, and for F_1_ hybrids between them (NY x GA only). Within pure NY and GA co-expression networks, we identified clusters of highly correlated genes, hereafter modules. We then tested for an effect of rearing temperature on module expression by the following method: (1) Expression for each module was summarized as the first principal component axis in a principal components analysis (hereafter module eigengene; (Langfelder and Horvath 2008)). (2) Rank-transformed module-eigengenes were used as response variables in a linear mixed effects model, which tested for the effect of rearing temperature on module expression. Individual fish length was used as a random covariate in the model to account for variation in size. *P*-values were corrected for multiple testing by False Discovery Rate correction (*Q*-value). Furthermore, we estimated the correlation between module eigengene and growth rate (mm per day) using Pearson correlation.

We compared pure GA and NY population networks to each other, and each to the hybrid network. Preservation analysis uses network-based preservation statistics to determine whether the properties of modules in a reference network can be identified in a test network. We used each pure population network as a reference, then tested whether modules in the other pure population, and the hybrid network, were preserved. In each comparison, the mean and variance of seven common network-based statistics are calculated using a permutation procedure, in which genes are randomly assigned to modules 200 times. This creates a null distribution of network statistics, against which *P*-values and *Z*-transformed scores for each statistic are calculated. A composite preservation statistic (*Z*_*summary*_) is calculated to summarize *Z*-scores across all preservation statistics following (Langfelder et al. 2011). Our interpretation of the results was based on guidelines by (Langfelder et al. 2011), in which simulated data suggest that *Z*_*summary*_ < 10 represents weak evidence of module preservation, while *Z*_*summary*_ < 2 suggests no preservation at all.

### Gene ontology analyses

To determine the functional role of each temperature-related and growth-correlated gene co-expression module in GA and NY, we performed gene ontology (GO) overrepresentation analysis using *topGO* in *R*. We used all expressed genes after filtering as the background gene list. GO terms with p-values below 0.05 were retained as significant.

### Impact of genetic differentiation and structural variants on regulatory dynamics

We tested if specific groups of genes (e.g. DEGs, inheritances classes, specific modules) are enriched inside one or multiple polymorphic inversions. First, we confirmed that known major inversions are polymorphic in our dataset by performing principal components analyses using *SNPrelate* (Zheng et al. 2012) based on SNPs within known inversions (Tigano et al. 2021; Akopyan et al. 2022). Inversion boundaries were defined based on linkage mapping data as described in the Supplementary text. We conservatively defined genes as being within an inversion if they started and ended within the particular inversion. For each inversion that was polymorphic in our study individuals (inversions on chr.4, chr.7, chr.8, chr.18(1,2,3), chr.19 and chr.24(2)) and each group of genes, we tested if the proportion of genes of a particular group is higher inside a particular inversion compared to the proportion of genes in the focal group across the collinear genome using Fisher’s Exact Tests in R.

## Author contributions

A.J., A.T., A.W., H.B., and N.O.T conceived of the project. Rearing (H.B., A.W.). Extraction (A.T.). Analysis (A.J., J.V.). First ms version (A.J., J.V.). All authors contributed to the final version.

## Acknowledgements

Thanks to Nicolas Lou, James Harrington and Chris Murray for assistance with rearing and sampling, and Harmony Borchardt-Wier for assistance with lab work. This study was funded through National Science Foundation grants to NOT (OCE-1756316) and HB (OCE-1756751)

## Data availability

All data are available on NCBI in the SRA under the BioProject PRJNA694674, with SRA accession SRR13523227 to SRR13523268.

## References

Akopyan M, Tigano A, Jacobs A, Wilder AP, Baumann H, Therkildsen NO. 2022. Comparative linkage mapping uncovers recombination suppression across massive chromosomal inversions associated with local adaptation in Atlantic silversides. Mol. Ecol. 00:1–19.

Barreto FS, Pereira RJ, Burton RS. 2015. Hybrid dysfunction and physiological compensation in gene expression. Mol. Biol. Evol. 32:613–622.

Berdan EL, Mérot C, Pavia H, Johannesson K, Wellenreuther M, Butlin RK. 2021. A large chromosomal inversion shapes gene expression in seaweed flies (Coelopa frigida). Evolution Letters 5:607–624.

Berner D. 2019. Allele Frequency Difference AFD -An Intuitive Alternative to FST for Quantifying Genetic Population Differentiation. Genes 10(4):308.

Billerbeck JM, Lankford TE Jr, Conover DO. 2001. Evolution of intrinsic growth and energy acquisition rates. I. Trade-offs with swimming performance in Menidia menidia. Evolution 55:1863–1872.

Bomblies K, Lempe J, Epple P, Warthmann N, Lanz C, Dangl JL, Weigel D. 2007. Autoimmune response as a mechanism for a Dobzhansky-Muller-type incompatibility syndrome in plants. PLoS Biol. 5:e236.

Bono LM, Smith LB Jr, Pfennig DW, Burch CL. 2017. The emergence of performance trade-offs during local adaptation: insights from experimental evolution. Mol. Ecol. 26:1720–1733.

Campbell-Staton SC, Velotta JP, Winchell KM. 2021. Selection on adaptive and maladaptive gene expression plasticity during thermal adaptation to urban heat islands. Nat. Commun. 12:1–14.

Cassone BJ, Molloy MJ, Cheng C, Tan JC, Hahn MW, Besansky NJ. 2011. Divergent transcriptional response to thermal stress by Anopheles gambiae larvae carrying alternative arrangements of inversion 2La. Mol. Ecol. 20:2567–2580.

Castel SE, Mohammadi P, Chung WK, Shen Y, Lappalainen T. 2016. Rare variant phasing and haplotypic expression from RNA sequencing with phASER. Nat. Commun. 7:12817.

Combs PA, Fraser HB. 2018. Spatially varying cis-regulatory divergence in Drosophila embryos elucidates cis-regulatory logic. PLoS Genet. 14:e1007631.

Conover DO, Clarke LM, Munch SB, Wagner GN. 2006. Spatial and temporal scales of adaptive divergence in marine fishes and the implications for conservation. J. Fish Biol. 69:21–47.

Conover DO, Duffy TA, Hice LA. 2009. The Covariance between Genetic and Environmental Influences across Ecological Gradients: Reassessing the Evolutionary Significance of Countergradient and Cogradient Variation. Ann. N. Y. Acad. Sci. 1168:100–129.

Conover DO, Heins SW. 1987. Adaptive variation in environmental and genetic sex determination in a fish. Nature 326:496–498.

Conover DO, Present TMC. 1990. Countergradient variation in growth rate: compensation for length of the growing season among Atlantic silversides from different latitudes. Oecologia 83:316–324.

Coolon JD, McManus CJ, Stevenson KR, Graveley BR, Wittkopp PJ. 2014. Tempo and mode of regulatory evolution in Drosophila. Genome Res. 24:797–808.

Crow T, Ta J, Nojoomi S, Aguilar-Rangel MR, Torres Rodríguez JV, Gates D, Rellán-Álvarez R, Sawers R, Runcie D. 2020. Gene regulatory effects of a large chromosomal inversion in highland maize. PLoS Genet. 16:e1009213.

Cutter AD, Bundus JD. 2020. Speciation and the developmental alarm clock. Elife 9:e56276

Dion-Côté A-M, Renaut S, Normandeau E, Bernatchez L. 2014. RNA-seq reveals transcriptomic shock involving transposable elements reactivation in hybrids of young lake whitefish species. Mol. Biol. Evol. 31:1188–1199.

Dobin A, Gingeras TR. 2015. Mapping RNA-seq Reads with STAR. Curr. Protoc. Bioinformatics 51:11.14.1–19.

Feder JL, Nosil P, Flaxman SM. 2014. Assessing when chromosomal rearrangements affect the dynamics of speciation: implications from computer simulations. Front. Genet. 5:295.

Filteau M, Pavey SA, St-Cyr J, Bernatchez L. 2013. Gene coexpression networks reveal key drivers of phenotypic divergence in lake whitefish. Mol. Biol. Evol. 30:1384–1396.

Findley AS, Monziani A, Richards AL, Rhodes K, Ward MC, Kalita CA, Alazizi A, Pazokitoroudi A, Sankararaman S, Wen X, et al. 2021. Functional dynamic genetic effects on gene regulation are specific to particular cell types and environmental conditions. Elife 10:e67077.

Fischer EK, Song Y, Hughes KA, Zhou W, Hoke KL. 2021. Nonparallel transcriptional divergence during parallel adaptation. Mol. Ecol. 30:1516–1530.

Fraser HB, Moses AM, Schadt EE. 2010. Evidence for widespread adaptive evolution of gene expression in budding yeast. Proc. Natl. Acad. Sci. U. S. A. 107:2977–2982.

Fuller ZL, Haynes GD, Richards S, Schaeffer SW. 2016. Genomics of Natural Populations: How Differentially Expressed Genes Shape the Evolution of Chromosomal Inversions in Drosophila pseudoobscura. Genetics 204:287–301.

Gould BA, Chen Y, Lowry DB. 2018. Gene regulatory divergence between locally adapted ecotypes in their native habitats. Mol. Ecol. 27:4174–4188.

Grishkevich V, Yanai I. 2013. The genomic determinants of genotype × environment interactions in gene expression. Trends Genet. 29:479–487.

Groszmann M, Greaves IK, Fujimoto R, Peacock WJ, Dennis ES. 2013. The role of epigenetics in hybrid vigour. Trends Genet. 29:684–690.

Han F, Jamsandekar M, Pettersson ME, Su L, Fuentes-Pardo AP, Davis BW, Bekkevold D, Berg F, Casini M, Dahle G, et al. 2020. Ecological adaptation in Atlantic herring is associated with large shifts in allele frequencies at hundreds of loci. Elife 9:e61076.

Hanson D, Hu J, Hendry AP, Barrett RDH. 2017. Heritable gene expression differences between lake and stream stickleback include both parallel and antiparallel components. Heredity 119:339–348.

Hart JC, Ellis NA, Eisen MB, Miller CT. 2018. Convergent evolution of gene expression in two high-toothed stickleback populations. PLoS Genet. 14:e1007443.

Hays CG, Hanley TC, Hughes AR, Truskey SB, Zerebecki RA, Sotka EE. 2021. Local Adaptation in Marine Foundation Species at Microgeographic Scales. Biol. Bull. 241:16–29.

Healy TM, Schulte PM. 2019. Patterns of alternative splicing in response to cold acclimation in fish. J. Exp. Biol. 222:jeb193516.

He G, He H, Deng XW. 2013. Epigenetic variations in plant hybrids and their potential roles in heterosis. J. Genet. Genomics 40:205–210.

Hice LA, Duffy TA, Munch SB, Conover DO. 2012. Spatial scale and divergent patterns of variation in adapted traits in the ocean. Ecol. Lett. 15:568–575.

Hu CK, York RA, Metz HC, Bedford NL, Fraser HB, Hoekstra HE. 2022. cis-Regulatory changes in locomotor genes are associated with the evolution of burrowing behavior. Cell Rep. 38:110360.

Hutchings JA, Swain DP, Rowe S, Eddington JD, Puvanendran V, Brown JA. 2007. Genetic variation in life-history reaction norms in a marine fish. Proc. Biol. Sci. 274:1693–1699.

Jacobs A, Elmer KR. 2021. Alternative splicing and gene expression play contrasting roles in the parallel phenotypic evolution of a salmonid fish. Mol. Ecol. 30:4955–4969.

Juneja P, Quinn A, Jiggins FM. 2016. Latitudinal clines in gene expression and cis-regulatory element variation in Drosophila melanogaster. BMC Genomics 17:981.

Kelley JL, Brown AP, Therkildsen NO, Foote AD. 2016. The life aquatic: advances in marine vertebrate genomics. Nat. Rev. Genet. 17:523–534.

Kirubakaran TG, Grove H, Kent MP, Sandve SR, Baranski M, Nome T, De Rosa MC, Righino B, Johansen T, Otterå H, et al. 2016. Two adjacent inversions maintain genomic differentiation between migratory and stationary ecotypes of Atlantic cod. Mol. Ecol. 25:2130–2143.

Kulmuni J, Westram AM. 2017. Intrinsic incompatibilities evolving as a by-product of divergent ecological selection: Considering them in empirical studies on divergence with gene flow. Mol. Ecol. 26:3093–3103.

Landry CR, Hartl DL, Ranz JM. 2007. Genome clashes in hybrids: insights from gene expression. Heredity 99:483–493.

Landry CR, Wittkopp PJ, Taubes CH, Ranz JM, Clark AG, Hartl DL. 2005. Compensatory cis-trans evolution and the dysregulation of gene expression in interspecific hybrids of Drosophila. Genetics 171:1813–1822.

Langfelder P, Horvath S. 2008. WGCNA: an R package for weighted correlation network analysis. BMC Bioinformatics 9:559.

Langfelder P, Luo R, Oldham MC, Horvath S. 2011. Is my network module preserved and reproducible? PLoS Comput. Biol. 7:e1001057.

Lankford TE Jr, Billerbeck JM, Conover DO. 2001. Evolution of intrinsic growth and energy acquisition rates. II. Trade-offs with vulnerability to predation in Menidia menidia. Evolution 55:1873–1881.

Lou RN, Fletcher NK, Wilder AP, Conover DO, Therkildsen NO, Searle JB. 2018. Full mitochondrial genome sequences reveal new insights about post-glacial expansion and regional phylogeographic structure in the Atlantic silverside (Menidia menidia). Mar. Biol. 165:124.

Love MI, Huber W, Anders S. 2014. Moderated estimation of fold change and dispersion for RNA-seq data with DESeq2. Genome Biol. 15:550.

Mach ME, Sbrocco EJ, Hice LA, Duffy TA, Conover DO, Barber PH. 2011. Regional differentiation and post-glacial expansion of the Atlantic silverside, Menidia menidia, an annual fish with high dispersal potential. Mar. Biol. 158:515–530.

Mack KL, Ballinger MA, Phifer-Rixey M, Nachman MW. 2018. Gene regulation underlies environmental adaptation in house mice. Genome Res. 28:1636–1645.

Mack KL, Campbell P, Nachman MW. 2016. Gene regulation and speciation in house mice. Genome Res. 26:451–461.

Mack KL, Nachman MW. 2017. Gene Regulation and Speciation. Trends Genet. 33:68–80.

Marquès-Bonet T, Cáceres M, Bertranpetit J, Preuss TM, Thomas JW, Navarro A. 2004. Chromosomal rearrangements and the genomic distribution of gene-expression divergence in humans and chimpanzees. Trends Genet. 20:524–529.

McGirr JA, Martin CH. 2020a. Ecological divergence in sympatry causes gene misexpression in hybrids. Mol. Ecol. 29:2707–2721.

McGirr JA, Martin CH. 2020b. Few fixed variants between trophic specialist pupfish species reveal candidate cis-regulatory alleles underlying rapid craniofacial divergence. Mol. Biol. Evol. 38:405–423.

McKenna A, Hanna M, Banks E, Sivachenko A, Cibulskis K, Kernytsky A, Garimella K, Altshuler D, Gabriel S, Daly M, et al. 2010. The Genome Analysis Toolkit: a MapReduce framework for analyzing next-generation DNA sequencing data. Genome Res. 20:1297–1303.

McManus CJ, Coolon JD, Duff MO, Eipper-Mains J, Graveley BR, Wittkopp PJ. 2010. Regulatory divergence in Drosophila revealed by mRNA-seq. Genome Res. 20:816–825.

Mérot C, Oomen RA, Tigano A, Wellenreuther M. 2020. A Roadmap for Understanding the Evolutionary Significance of Structural Genomic Variation. Trends Ecol. Evol. 35:561–572.

Moran BM, Payne C, Langdon Q, Powell DL, Brandvain Y, Schumer M. 2021. The genomic consequences of hybridization. Elife 10:e69016.

Mugal CF, Wang M, Backström N, Wheatcroft D, ålund M, Sémon M, McFarlane SE, Dutoit L, Qvarnström A, Ellegren H. 2020. Tissue-specific patterns of regulatory changes underlying gene expression differences among Ficedula flycatchers and their naturally occurring F1 hybrids. Genome Res. 30:1727–1739.

Naseeb S, Carter Z, Minnis D, Donaldson I, Zeef L, Delneri D. 2016. Widespread Impact of Chromosomal Inversions on Gene Expression Uncovers Robustness via Phenotypic Buffering. Mol. Biol. Evol. 33:1679–1696.

Navarro A, Barton NH. 2003. Accumulating postzygotic isolation genes in parapatry: a new twist on chromosomal speciation. Evolution 57:447–459.

Oomen RA, Juliussen E, Olsen EM, Knutsen H, Jentoft S, Hutchings JA. 2021. Cryptic microgeographic variation in responses of larval Atlantic cod to warmer temperatures. bioRxiv. doi: 10.1101/2021.02.03.429645v1

Ortíz-Barrientos D, Counterman BA, Noor MAF. 2007. Gene expression divergence and the origin of hybrid dysfunctions. Genetica 129:71–81.

Pavey SA, Collin H, Nosil P, Rogers SM. 2010. The role of gene expression in ecological speciation. Ann. N. Y. Acad. Sci. 1206:110–129.

Payne C, Bovio S, Powell D, Gunn T, Banerjee S, Grant V, Rosenthal G, Schumer M. 2021. Genomic insights into variation in thermotolerance between hybridizing swordtail fishes. bioRxiv. doi: 10.1101/2021.10.22.465484v1

Pereira RJ, Barreto FS, Burton RS. 2014. Ecological novelty by hybridization: experimental evidence for increased thermal tolerance by transgressive segregation in Tigriopus californicus. Evolution 68:204–215.

Pettersson ME, Rochus CM, Han F, Chen J, Hill J, Wallerman O, Fan G, Hong X, Xu Q, Zhang H, et al. 2019. A chromosome-level assembly of the Atlantic herring genome-detection of a supergene and other signals of selection. Genome Res. 29:1919–1928.

Renaut S, Nolte AW, Bernatchez L. 2009. Gene expression divergence and hybrid misexpression between lake whitefish species pairs (Coregonus spp. Salmonidae). Mol. Biol. Evol. 26:925–936.

Said I, Byrne A, Serrano V, Cardeno C, Vollmers C, Corbett-Detig R. 2018. Linked genetic variation and not genome structure causes widespread differential expression associated with chromosomal inversions. Proc. Natl. Acad. Sci. U. S. A. 115:5492–5497.

Sanford E, Kelly MW. 2011. Local adaptation in marine invertebrates. Ann. Rev. Mar. Sci. 3:509–535.

Signor SA, Nuzhdin SV. 2018. The Evolution of Gene Expression in cis and trans. Trends Genet. 34:532–544.

Sodeland M, Jorde PE, Lien S, Jentoft S, Berg PR, Grove H, Kent MP, Arnyasi M, Olsen EM, Knutsen H. 2016. “Islands of Divergence” in the Atlantic Cod Genome Represent Polymorphic Chromosomal Rearrangements. Genome Biol. Evol. 8:1012–1022.

Stacklies W, Redestig H, Scholz M, Walther D, Selbig J. 2007. pcaMethods—a bioconductor package providing PCA methods for incomplete data. Bioinformatics 23:1164–1167.

Stevison LS, Hoehn KB, Noor MAF. 2011. Effects of inversions on within-and between-species recombination and divergence. Genome Biol. Evol. 3:830–841.

Therkildsen NO, Wilder AP, Conover DO, Munch SB, Baumann H, Palumbi SR. 2019. Contrasting genomic shifts underlie parallel phenotypic evolution in response to fishing. Science 365:487–490.

Tigano A, Jacobs A, Wilder AP, Nand A, Zhan Y, Dekker J, Therkildsen NO. 2021. Chromosome-Level Assembly of the Atlantic Silverside Genome Reveals Extreme Levels of Sequence Diversity and Structural Genetic Variation. Genome Biol. Evol. 13:evab098.

Velotta JP, Wegrzyn JL, Ginzburg S, Kang L, Czesny S, O’Neill RJ, McCormick SD, Michalak P, Schultz ET. 2017. Transcriptomic imprints of adaptation to freshwater: parallel evolution of osmoregulatory gene expression in the Alewife. Mol. Ecol. 26:831–848.

Verta J-P, Jacobs A. 2022. The role of alternative splicing in adaptation and evolution. Trends Ecol. Evol. 37(4):299–308

Verta J-P, Jones FC. 2019. Predominance of cis-regulatory changes in parallel expression divergence of sticklebacks. Elife 8:e43785.

Wadgymar SM, Lowry DB, Gould BA, Byron CN, Mactavish RM, Anderson JT. 2017. Identifying targets and agents of selection: innovative methods to evaluate the processes that contribute to local adaptation. Methods Ecol. Evol. 8:738–749.

Wang L, Israel JW, Edgar A, Raff RA, Raff EC, Byrne M, Wray GA. 2020. Genetic basis for divergence in developmental gene expression in two closely related sea urchins. Nat Ecol Evol 4:831–840.

Wilder AP, Palumbi SR, Conover DO, Therkildsen NO. 2020. Footprints of local adaptation span hundreds of linked genes in the Atlantic silverside genome. Evolution Letters 4:430–443.

Wittkopp PJ, Haerum BK, Clark AG. 2004. Evolutionary changes in cis and trans gene regulation. Nature 430:85–88.

Wittkopp PJ, Kalay G. 2011. Cis-regulatory elements: molecular mechanisms and evolutionary processes underlying divergence. Nat. Rev. Genet. 13:59–69.

Wray GA, Hahn MW, Abouheif E, Balhoff JP, Pizer M, Rockman MV, Romano LA. 2003. The evolution of transcriptional regulation in eukaryotes. Mol. Biol. Evol. 20:1377–1419.

Yamahira K, Conover DO. 2002. INTRA-VS. INTERSPECIFIC LATITUDINAL VARIATION IN GROWTH: ADAPTATION TO TEMPERATURE OR SEASONALITY? Ecology 83:1252–1262.

York RA, Patil C, Abdilleh K, Johnson ZV, Conte MA, Genner MJ, McGrath PT, Fraser HB, Fernald RD, Streelman JT. 2018. Behavior-dependent cis regulation reveals genes and pathways associated with bower building in cichlid fishes. Proc. Natl. Acad. Sci. U. S. A. 115:E11081–E11090.

Zheng X, Levine D, Shen J, Gogarten SM, Laurie C, Weir BS. 2012. A high-performance computing toolset for relatedness and principal component analysis of SNP data. Bioinformatics 28:3326–3328.

