## Supplementary Material for "Temperature-dependent gene regulatory divergence underlies local adaptation with gene flow in the Atlantic silverside"

#### Supplementary methods

**Linkage mapping.** To improve the contiguity of the Atlantic silverside reference genome (Tigano et al. 2021), we anchored the genome assembly to a RAD-seq based female Georgia linkage map (Akopyan et al. 2022). However, the linkage map used for this assembly differs marginally from the one in (Akopyan et al. 2022), as it was constructed from genome-aligned RAD-seq data rather than *de novo* assembled loci Overall, both linkage maps are highly comparable. The linkage map was constructed as described in (Akopyan et al. 2022) with slight modifications outlined here:

In brief, we used ddRAD sequencing (Peterson et al. 2012) to identify and genotype single nucleotide polymorphisms (SNPs) for linkage map construction from 568 individuals across five families, including the two founders, 138 F1 offspring, six additional $F_1$ siblings and their 282 $F_2$ offspring. Reads were processed in Stacks v1.48 (Catchen et al. 2013) with the module *process_radtags* to discard low-quality reads and reads with ambiguous barcodes or RAD cut-sites. The remaining reads were demultiplexed and aligned to the *Menidia menidia* reference genome v1 (Tigano et al. 2021) using Bowtie2 v2.2.9 (-*very-sensitive*). We only retained those reads that were uniquely mapped to the reference genome and extracted RAD loci with: i) minimum read depth of three, ii) minimum mapping quality of 10, iii) and maximum clipped proportion of 0.15. Variant calling was also performed with *pstacks* using the default SNP model with a genotype likelihood ratio test critical value ($\alpha$) of 0.05. We built a catalog of all loci using parents (and grandparents for the $F_2$ generation) with *cstacks*, and matched progeny against the catalog using *sstacks*. The *populations* module was used to filter variants to retain only the first SNP per locus and generated a VCF file for each of the two $F_1$ families, and one for the $F_2$ generation including the three intercross families.

We constructed one female linkage map for each of our three crosses (F1 GAxNY, F1 NYxGA, F2) using *Lep-MAP3* (Rastas 2017). In brief, offspring genotypes were called by accounting genotype information of parents (and grandparents in $F_2$ family) with the

*ParentCall2* module, markers with high segregation distortion were removed using the distortionLod=1 option in *SeparateChromosomes2*, separated markers were merged into linkage groups with a logarithm of odds (LOD) score limit of 20 and minimum linkage group size of 10 markers using markers informative in females only, and we used the *OrderMarkers2* module to compute genetic distances in centimorgan (i.e., recombination rates) between all adjacent markers for each linkage group using the default Haldane's mapping function. We used maternally informative markers to construct the $F_1$ maps, and both maternally and dually informative markers to construct the $F_2$ map.

We used the female $F_1$ linkage map for the Georgia population to anchor and order the Atlantic silverside reference genome v1 scaffolds into chromosomes using *AllMaps* (Tang et al. 2015). Chromosomes were renamed based on synteny with the medaka genome. Furthermore, we converted the coordinates of RAD loci for all linkage maps from scaffold to anchored chromosome coordinates using *CrossMap v.0.1.4* (Zhao et al. 2014) and identified inversions between NY and GA by comparing the F1 NY-linkage map and F2 linkage map to the GA-anchored reference genome (see (Akopyan et al. 2022) for details).

Lastly, we re-annotated the anchored genome assembly (*M. menidia* reference genome v2) following the pipeline outlined in (Tigano et al. 2021). In brief, first we identified and annotated repeats using *Repeatmodeler2* (Flynn et al. 2020) and *Repeatmasker* (Smit et al. 2015). Next, we used *BRAKER2* (Brůna et al. 2021) with evidence from: i) RNA-seq for diverse Atlantic silverside individuals from different populations and at different developmental stages (Tigano et al. 2021; Therkildsen and Baumann 2020); ii) protein-homology evidence from six different teleost species and the UniProtKB (Swiss-prot) database. Subsequently, we performed five iterative rounds in *MAKER* as described in (Tigano et al. 2021). Lastly, functionally annotated the predicted genes using Blast2GO in Omnibox v.1.2.4 (Gotz et al. 2008) using the UniProtKB (Swiss-Prot) database and InterProScan2 (Zdobnov and Apweiler 2001).

**Criteria for inheritance modes.** We identified the inheritance mode for each gene based on the following criteria outlined in (Coolon et al. 2014):

- *conserved:* same expression in both parental populations and the F1 hybrids.
- *NY dominant:* Expression in F1 hybrids is similar to NY, but different from GA.
- *GA dominant*: Expression in F1 hybrids is similar to GA, but different from NY.
- *Additive*: Expression in F1 hybrids differs from both parental populations, but is intermediate in expression between both parental populations.
- *(Transgressive) Overdominant*: Expression in F1 hybrids is higher than in both parental populations.
- *(Transgressive) Underdominant*: Expression in F1 hybrids is lower than in both parental populations.

- *Ambiguous*: Expression variation does not clearly fit one of the patterns above.

**Criteria for regulatory modes.** We identified the regulatory mode for each gene based on the following criteria outlined in (Coolon et al. 2014):

- *all-cis: cis*-differences alone explain expression differences between populations
- *all-trans*: *trans*-differences alone explain expression differences between populations
- *cis + trans*: additive *cis* and *trans*-effects explain expression differences between populations (*cis* and trans can either act in the same or opposite directions)
- *cis x trans*: *cis*- and *trans*-effects that partially cancel each other out
- *compensatory*: *cis*- and *trans*-effects of equal and opposite sign. No difference in expression between parental populations.
- *Conserved*: No regulatory difference.
- *Ambiguou*s: not conserved and not statistically supported as one of the previous classifications
- *Uninformative*: Genes with low expression counts (<3) in either parent or hybrids.

### Supplementary results

**Comparisons of growth rates:** Comparing growth rates (mm per day) between groups and temperatures showed a significant difference between populations (ANOVA: $F_{Group} = 81.246$, $p = 4.51e-14$; df = 2), with NY individuals growing significantly faster than GA individuals (Fig. 1b). In both populations, growth rates differed between temperatures (ANOVA: $F_{Temperature} = 210.767$, $p < 2e-16$; df = 1; $F=_{(Group*Temperature)} = 1.346$, $p = 0.273$, df =2), with faster growth at 26ºC with evidence for additive inheritance at 20ºC ($p < 0.05$) but NY-dominance and/or maternal effects at 26ºC, as growth rate of the HYB group was not different from NY individuals ($p = 0.12$) but significantly different from GA ($p < 0.001$). This might indicate a significant effect of the environment on potential outcomes of hybridization.

**Linkage mapping and genome anchoring:**
In total, we obtained 2.13 billion ddRADseq reads (~3.3 million per individual), of which 10% of which were removed after filtering with Stacks, and after mapping to the genome ~2 million uniquely mapped reads were retained per individual. Following genotype calling and filtering, we retained 50,955 SNPs with on average 15.5x depth of coverage across individuals in the Georgia $F_1$ family and 49,115 SNPs at on average 16.3x in the New York $F_1$ family, and 71,612 SNPs at on average 13x coverage in the $F_2$ family. After further filtering in *Lep-Map3*, we constructed linkage maps with 9,337 and 9,769 maternally informative SNPs in the Georgia and New York $F_1$ families respectively, and 18,651 maternally and dually informative SNPs in

the $F_2$ family. For each of the three genetic maps, we obtained 24 linkage groups, consistent with the haploid number of *M. menidia* chromosomes (*n*=24, (Warkentine et al. 1987)). The Georgia female map spans 3667.5 cM, the New York female map spans 3198.6 cM, and the $F_2$ map, which contains both maternally and dually informative markers, spans 3490.5 cM.

We were able to anchor 74.6 % of the genome into 24 chromosomes (186 scaffolds; Table S8-9). Comparing the NY F1 linkage map to the reference genome from a Georgia-origin individual revealed eight inversions on six chromosomes (Akopyan et al. 2022). The inversions ranged in size from 0.7-11.9 megabases (Mb), with the largest spanning much of the length of chromosome 8. The majority of chromosomes 18 and 24 are also inverted, with the former having three adjacent inversions at positions 4.2-7.9 Mb, 9.3-11.0 Mb, and 11.1-11.8 Mb (treated as a single inversion in this study), and the latter having the second largest inversion that captures 8.8 Mb. Smaller inversions are seen on chromosome 4 (at position 12.2-14.1 Mb), chromosome 7 (at position 5.9-7.6 Mb), and chromosome 19 (at position 2.2-3.4 Mb).

**Across temperature differential expression:** We detected 4119 differentially expressed genes (DEGs) (22.2% of all expressed genes) between NY and GA ($q < 0.05$), when individuals were combined across both temperatures. This number is lower than the sum of DE genes from both temperatures separately, as temperature-sensitive genes that show different patterns between the two temperatures will likely not be detected as DE between NY and GA. Due to variation in total length within and between groups (supplementary fig. 1), we also tested for genes with expression patterns associated with total length, but no gene was significantly associated with length ($q < 0.05$), suggesting that length variation has no or only a negligible effect on our results. Comparing hybrids to the maternal and paternal populations across temperatures, we detected 3306 DEGs between hybrids and the maternal NY population and 6615 DEGs between hybrids and the paternal GA population. The larger number of DEGs between hybrids and the paternal population (49.98% more DEGs) compared to the maternal population supports the presence of significant maternal effects on gene expression.

**Co-evolution of *cis*- and *trans*-regulation:** We both detected genes showing compensatory evolution, where *trans* and *cis* factors have equal but opposite effects (*compensatory*), partially canceling each other out (*cis x trans*), and genes where regulatory mechanisms are additive and reinforce each other (*cis + trans*). Slightly more genes showed evidence for compensatory evolution at 20°C (n=57) compared to 26°C (n=39) (Fig. S5). Only one and three genes showed evidence for reinforcing interactions (*cis + trans*) at 20°C and 26°C, respectively, which is indicative of directional selection. However, we found that nine genes that showed *all-trans* regulation under one temperature, showed an interaction of *cis*- and *trans*-effects under the other temperature, with two genes showing reinforcing *cis + trans* regulation, seven genes *cis x trans* interaction, and another seven showing fully compensatory regulation. However, overall, only a small number of genes were available for allele-specific expression analysis, potentially reducing our ability to draw more general conclusions.

**Regulatory changes of Hub genes.** To determine the impact of regulatory changes at individual genes on gene network properties, we compared the expression, regulatory modes and inheritance of 'Hub genes', the most strongly connected genes in each co-expression module, within and between populations and temperature regimes.

We found that Hub genes, the top 5% connected genes, in both networks were significantly enriched for DE genes between NY and GA, with 284 (HGT: p = 1.161011e-06) and 299 (HGT: p = 4.117635e-09) of the 938 Hub genes being differentially expressed at 20ºC in the GA and NY networks, respectively. At 26ºC, 447 (HGT: p = 4.257295e-48) and 441 (HGT: p = 8.709487e-34) of the 938 hub were differentially expressed between NY and GA in the GA and NY networks, respectively. The greater enrichment at 26ºC is in line with the higher number of DE genes at 26º compared to 20ºC (Fig. 2A). A significant proportion of Hub genes in both the NY and GA networks were misexpressed in hybrids at 20ºC (HGT; GA: p= 8.9e-18; NY: p = 1.6e-37). Although there are more differentially expressed Hub genes at 26ºC, we did not detect an enrichment of misexpressed genes at that temperature, likely due to the generally low number of misexpressed genes at 26ºC.

While we detected regulatory divergence of some Hub genes, e.g. one *all-cis* regulated Hub gene in each network at both temperatures, representing between 2% and 3% of all tested Hub genes, neither *all-trans* nor *all-cis* regulated genes were statistically overrepresented at either temperature (HGT: p > 0.05). Interestingly, while the same Hub gene was *all-cis* divergent at both temperatures in the NY network (*SODM*), a different Hub gene showed all-cis divergent between temperatures in the GA network (uncharacterized locus at 20ºC; *OGA* at 26ºC), supporting the importance of temperature-dependent gene regulatory architectures. Most genes with evidence for regulatory divergence, were *all-trans* regulated, with a few other hub genes that are *cis x trans* or *compensatory* regulated. Overall, the majority of genes showed ambiguous, conserved or uninformative regulatory modes.

**Supplementary figures:**

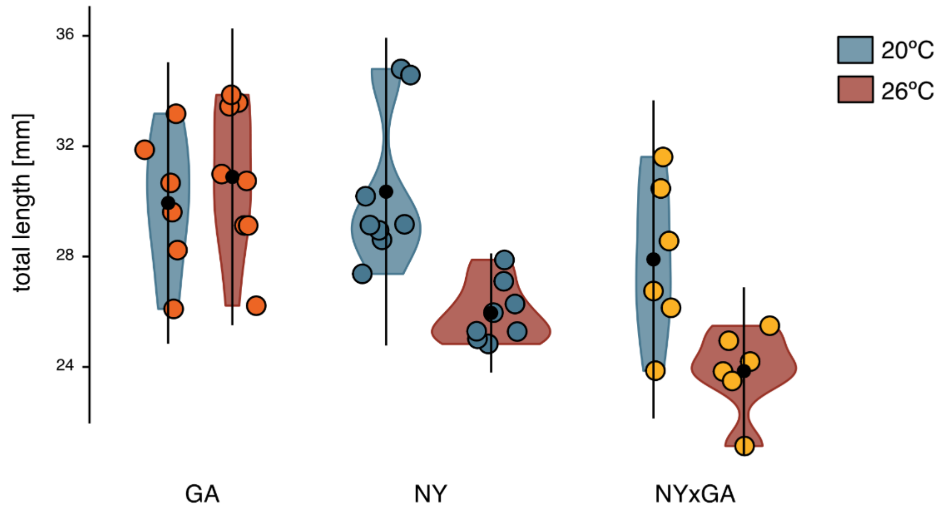


**Fig S1. Body length distribution of all individuals used for RNAseq.** Total length in mm at time of sampling is plotted for all individuals by cross and rearing temperature, with coloured points showing individuals. The black dot shows the mean ± standard deviation (black line), and the violin plots show the distribution of values for each group.

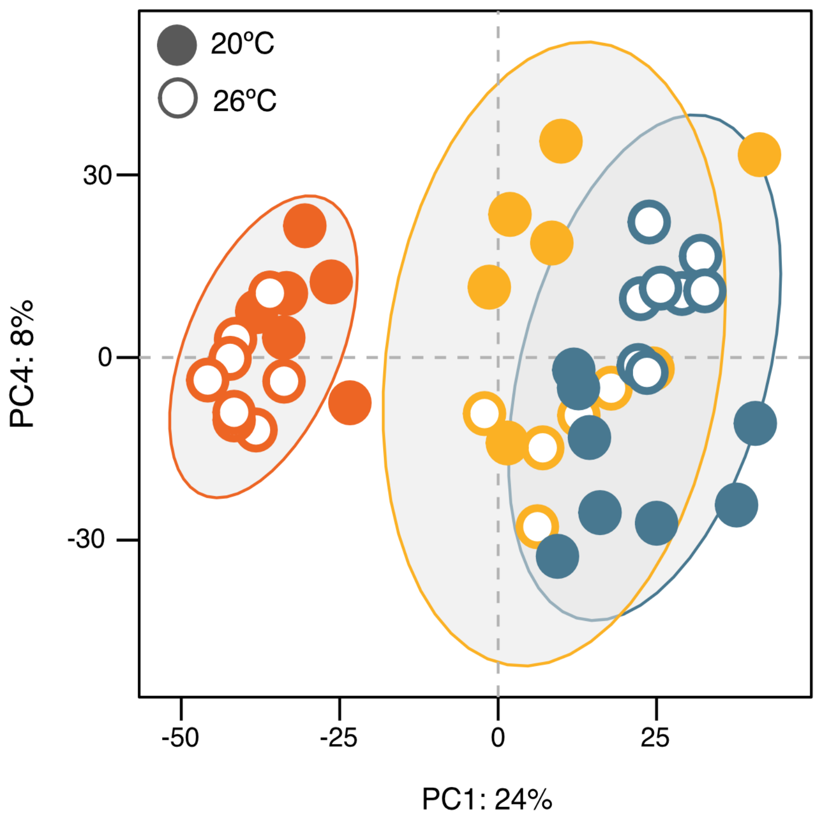


**Fig S2. Gene expression PCA.** Principal components 1 and 4 based on gene expression read counts. GA individuals are shown in orange-red, NY individuals in blue, and NYxGA hybrids in yellow.

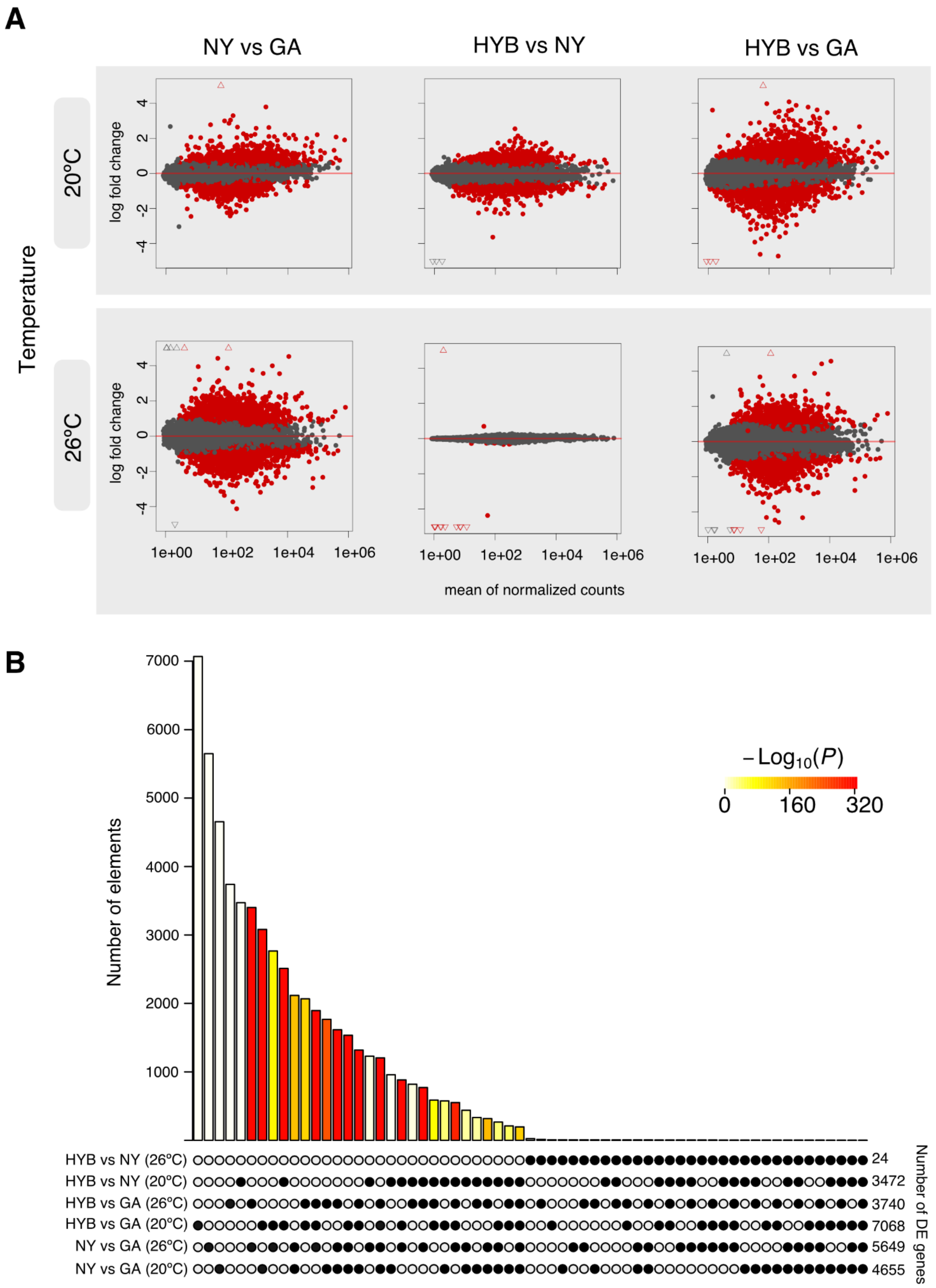


**Fig S3. Differential gene expression. A)** MA plots showing differential gene expression between groups by temperature regime. Significantly differentially expressed genes are highlighted in red. Genes with log-fold changes outside the plotted range are shown as triangles. **B)** Overlap of differentially expressed genes between comparisons. The barplot shows the number of differentially expressed (DE)

genes per group and per comparison. The black dots below show which group or comparison each bar corresponds to. Colours show the $-\log_{10}$(p-values) from Fisher's Exact Tests for each comparison.

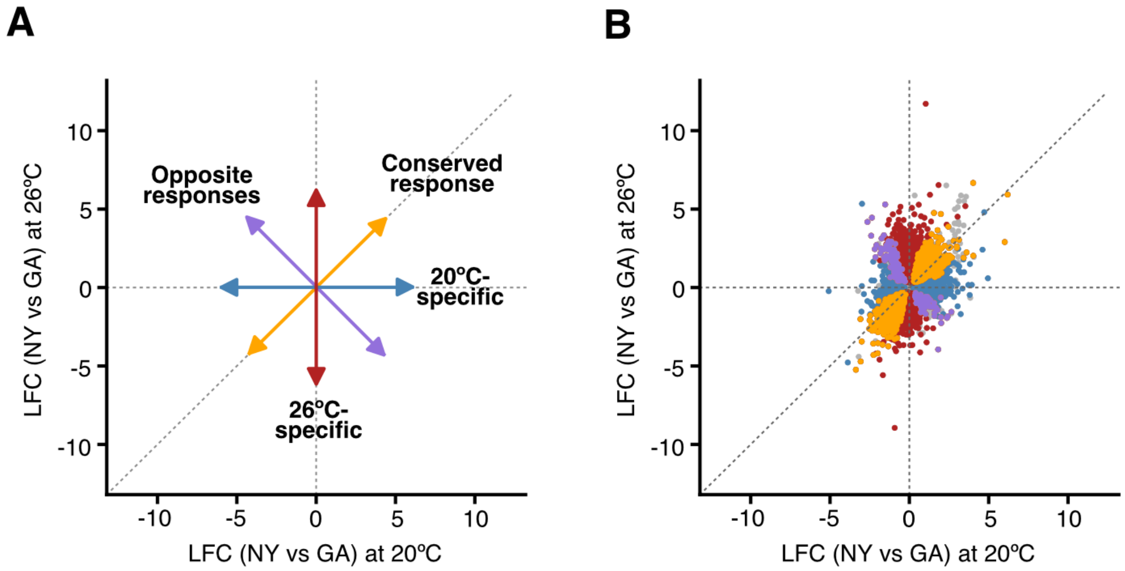


**Fig S4. Temperature-dependent gene expression divergence between NY and GA. A)** Theoretical patterns for temperature-dependent expression divergences. **B)** Comparison of empirical expression divergences between temperature-regimes. Individual genes (points) are coloured depending on their expression pattern between the two temperature regimes as indicated in panel A. Non-differentially expressed genes (adjusted p-value < 0.05) are in grey. The x- and y-axis show the log-fold change (LFC) for the NY vs GA comparison at 20° and 26°C, respectively.

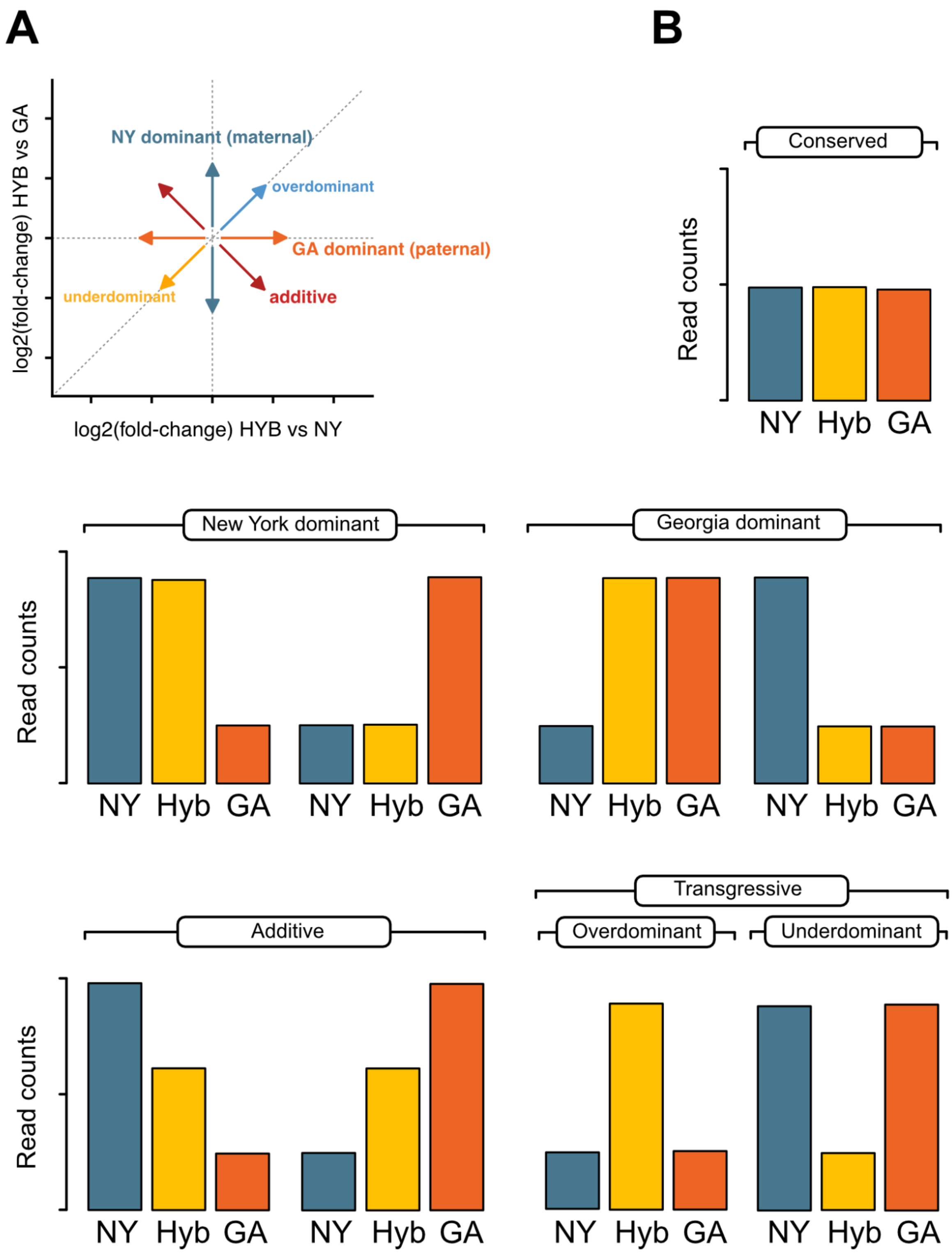


**Fig S5. Illustration of gene expression inheritance modes. A)** Gene expression modes are inferred based on the comparison of gene expression between both parental populations and each parental population and the hybrids. Inheritance modes are explained in the Supplementary methods. **B)** Theoretical expression patterns for the different inheritance modes. The height of the barplot represents the hypothetical expression of a single gene in the different populations.

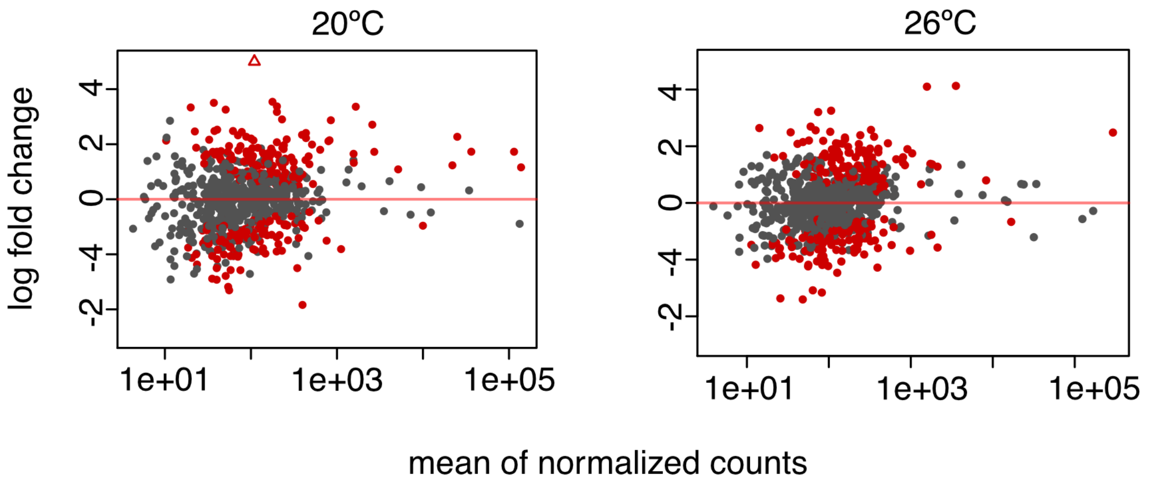


**Fig S6. Allele-specific expression (ASE).** MA plots showing log-fold change between NY and GA alleles at 20°C and 26°C. Genes showing significant ASE are highlighted in red. Genes with log-fold changes outside the plotted range are shown as triangles.

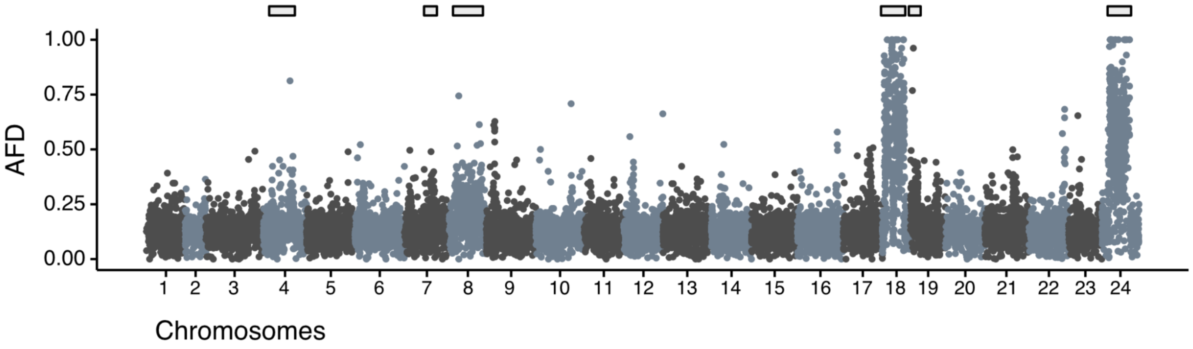


**Fig. S7. Manhattan plot** of Allele Frequency Differentiation (AFD) between individuals from GA and NY based on SNPs called from RNA-seq data. The locations of inversion segregating between populations are highlighted by grey boxes.

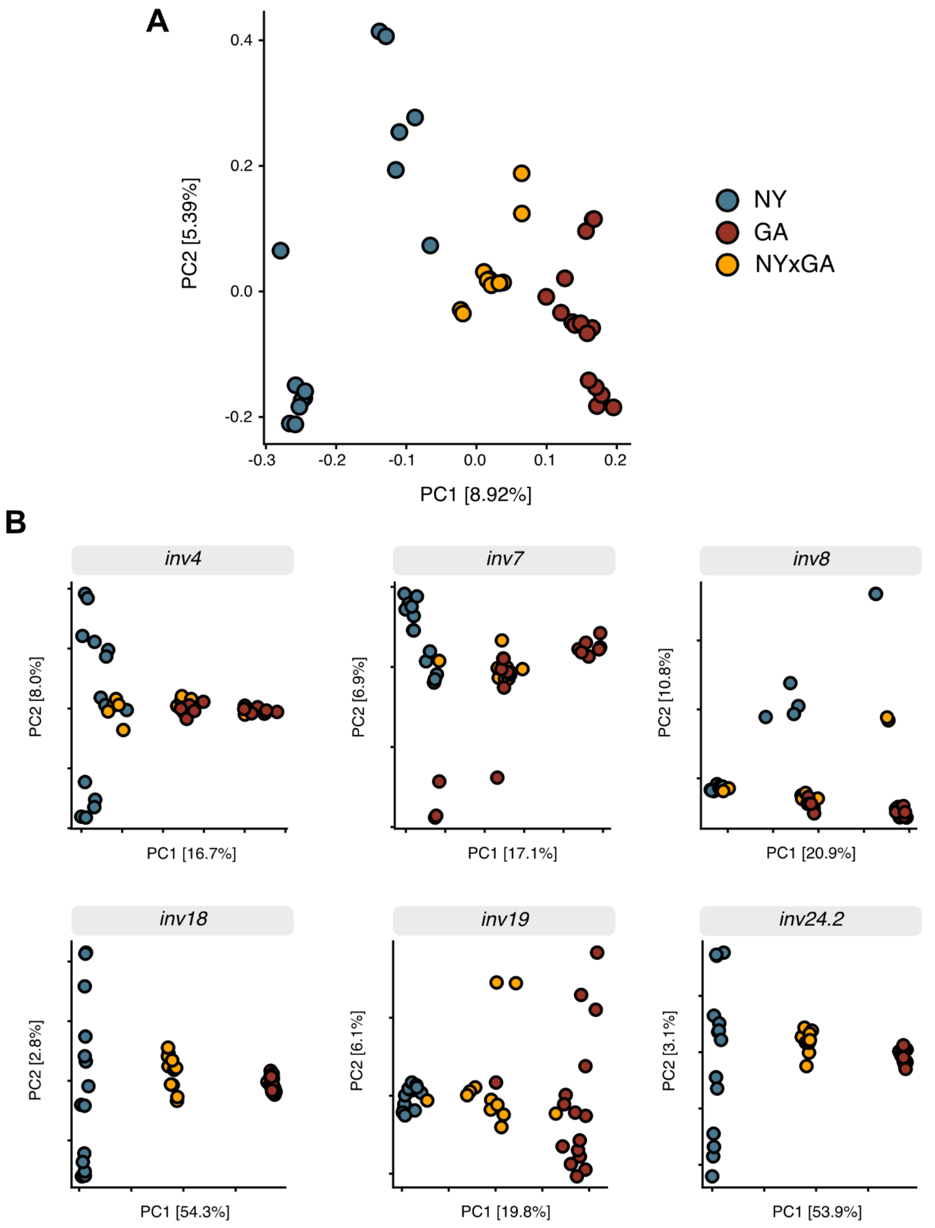


**Fig S8. Principal components analysis. A)** PCA for all chromosomes based on an LD-pruned SNP dataset. **B)** PCAs for SNPs within each inversion region segregating between our study populations. F1 hybrids are not always heterozygous for all inversions as some parents used in the crosses were likely not homozygous for alternative karyotypes.

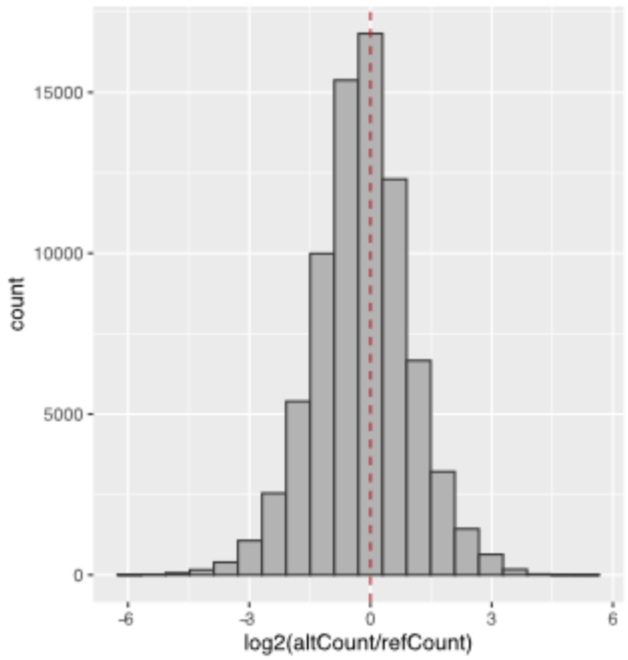


**Fig S9. ASE read count histogram.** Distribution of read count ratios between the NY and GA alleles.

**Supplementary tables:**

**Table S1. Sample information.** Sample number and average body length for individuals from New York (NY), Georgia (GA) and their F1 hybrids (NY x GA) reared at two different temperatures (20°C and 26°C).

| Population/ Group | Temperature | N. Individuals | Hatching (dpf) | Sampling age (dpf) | Mean size (mm) ± s.d. |
|---|---|---|---|---|---|
| NY | 20°C | 8 | 9 | 58-65 | 30.22 ± 2.79 |
| NY | 26°C | 8 | 5 | 37 | 25.96 ± 1.08 |
| GA | 20°C | 6 | 9 | 87 | 29.94 ± 2.55 |
| GA | 26°C | 8 | 5 | 58 | 30.89 ± 2.69 |
| NY x GA | 20°C | 6 | 9 | 65 | 27.90 ± 2.88 |
| NY x GA | 26°C | 6 | 5 | 37-45 | 23.85 ± 1.52 |

**Table S2. Differential expression in population comparisons.** Numbers of differentially expressed (DE) genes by comparison between individuals from New York (NY), Georgia (GA) and their F1 hybrids (HYB - NY mother/GA father).

| Group1 | Group2 | N. DE genes (FDR < 0.05) | Upregulated in Group1 | Downregulated in Group1 |
|---|---|---|---|---|
| NY at 20°C | GA at 20°C | 4655 | 2674 | 1981 |
| NY at 26°C | GA at 26°C | 5649 | 2857 | 2792 |
| HYB at 20°C | NY at 20°C | 3472 | 1790 | 1682 |
| HYB at 20°C | GA at 20°C | 7068 | 3519 | 3549 |
| HYB at 26°C | NY at 26°C | 24 | 2 | 22 |
| HYB at 26°C | GA at 26°C | 3740 | 1775 | 1965 |

**Table S3. GxE, thermal plasticity.** Number of DE genes between rearing temperature within each group: New York (NY), Georgia (GA) and NYxGA F1 hybrids (HYB).

| 20°C | 26°C | N. DE genes (FDR < 0.05) | Upregulated at 20°C | Downregulated at 20°C |
|---|---|---|---|---|
| NY | NY | 1504 | 1005 | 499 |
| GA | GA | 2371 | 1310 | 1061 |
| HYB | HYB | 166 | 38 | 128 |

**Table S4.** Number of genes by inheritance mode at 20°C and 26°C**.** Expression patterns for each inheritance mode are shown in Fig. S5 and explained in the supplementary methods.

| **Inheritance mode** | **N. genes at 20°C (proportion of all expressed genes)** | **N. genes at 26°C (proportion of all expressed genes)** |
|---|---:|---:|
| Conserved | 8281 (44.8%) | 11098 (60.0%) |
| Additive | 6 (0.03%) | 0 (0.0%) |
| NY dominant | 2498 (13.5%) | 3397 (18.4%) |
| GA dominant | 382 (2.1%) | 0 (0.0%) |
| Overdominant | 1393 (7.5%) | 1 (0.005%) |
| Underdominant | 1112 (6.0%) | 12 (0.07%) |
| Ambiguous | 4823 (26.2%) | 3974 (21.7%) |

**Table S5.** Number of genes by regulatory mode. We were able to infer regulatory modes for X genes at 20°C and X genes at 26°C**.**

| **Regulatory mode** | **N. genes at 20°C** | **N. genes at 26°C** |
|---|---|---|
| Conserved | 143 | 192 |
| Compensatory | 28 | 18 |
| cis x trans | 19 | 15 |
| cis + trans | 2 | 4 |
| all-cis | 21 | 16 |
| all-trans | 52 | 102 |
| ambiguous | 130 | 209 |
| Uninformative | 224 | 88 |

**Table S6.** New York (NY) co-expression network modules and corresponding summary statistics for ANCOVA on the effect of temperature acclimation on module eigengene, and correlation analysis between module eigengene and growth rate (mm per day). $Z_{summary}$ statistics for module preservation analysis for each comparison (NY vs. GA, NY vs. Hybrid) are also presented. $Z_{summary}$ scores < 10 are considered weak support for between-network preservation. P-values in bold are significant after false-discovery rate correction for multiple testing.

| | | Temp ANCOVA | | Growth correlation | | Module preservation | |
|---|---|---|---|---|---|---|---|
| **Module** | **Size** | **F** | **P** | **r** | **P** | **$Z_{summary}$ NYvGA** | **$Z_{summary}$ NYvHYB** |
| black | 715 | 0.2 | 0.8 | -0.15 | 0.7 | 28.8 | 41.2 |
| blue | 2735 | 0.3 | 0.8 | -0.29 | 0.4 | 68.2 | 52.3 |
| brown | 1484 | 45.6 | **<0.001** | 0.93 | **<0.001** | 5.1 | 12.5 |
| cyan | 299 | 1.1 | 0.5 | -0.47 | 0.1 | 10.1 | 8.6 |
| darkgreen | 66 | 7.1 | **0.05** | 0.70 | 0.01 | 2.4 | 7.4 |
| darkred | 98 | 0.1 | 0.8 | -0.15 | 0.7 | 24.6 | 8.8 |
| darkturquoise | 43 | 6.2 | **0.05** | 0.37 | 0.3 | 10.3 | 8.6 |
| green | 1329 | 30.5 | **0.001** | 0.56 | 0.06 | 12.2 | 3.7 |
| greenyellow | 443 | 43.8 | **<0.001** | 0.75 | **0.006** | 15.6 | 10.0 |
| grey60 | 212 | 16.0 | **0.006** | 0.57 | 0.06 | 4.4 | 7.3 |
| lightcyan | 227 | 0.0 | 0.8 | 0.23 | 0.61 | 10.1 | 9.2 |
| lightgreen | 137 | 0.1 | 0.8 | -0.10 | 0.8 | 4.8 | 2.6 |
| lightyellow | 125 | 1.0 | 0.5 | 0.11 | 0.8 | 3.7 | 4.4 |
| magenta | 577 | 44.4 | **<0.001** | -0.78 | **<0.001** | 16.8 | 22.0 |
| midnightblue | 253 | 0.5 | 0.7 | -0.21 | 0.6 | 7.6 | 4.5 |
| pink | 632 | 0.4 | 0.8 | 0.11 | 0.9 | 59.2 | 23.3 |
| purple | 542 | 5.0 | 0.08 | 0.52 | 0.09 | 14.1 | 22.8 |
| red | 1085 | 6.4 | 0.05 | 0.53 | 0.09 | 19.1 | 5.2 |
| royalblue | 116 | 0.1 | 0.8 | -0.01 | 1 | 4.3 | 5.8 |

| | | | | | | | |
|---|---|---|---|---|---|---|---|
| salmon | 325 | 10.<br>7 | **0.02** | 0.65 | **0.02** | 10.5 | 5.5 |
| tan | 335 | 8.1 | **0.04** | -0.29 | 0.4 | 15.0 | 10.8 |
| turquoise | 3961 | 45.<br>2 | **<0.001** | -0.77 | **0.006** | 19.9 | 10.5 |
| yellow | 1462 | 10.<br>3 | **0.02** | 0.58 | 0.06 | 13.0 | 19.5 |

**Table S7.** Georgia (GA) co-expression network modules and corresponding summary statistics for ANCOVA on the effect of temperature acclimation on module eigengene, and correlation analysis between module eigengene and growth rate (mm per day). $Z_{summary}$ statistics for module preservation analysis for each comparison (GA vs. NY, GA vs. Hybrid) are also presented. $Z_{summary}$ scores < 10 are considered weak support for between-network preservation. P-values in bold are significant after false-discovery rate correction for multiple testing.

| | | **Temp ANCOVA** | | **Growth correlation** | | **Module preservation** | |
|---|---|---|---|---|---|---|---|
| **Module** | **Size** | **F** | **P** | **r** | **P** | **$Z_{summary}$ NYvGA** | **$Z_{summary}$ NYvHYB** |
| black | 883 | 8.4 | 0.07 | 0.62 | 0.09 | 8.3 | 9.4 |
| blue | 2405 | 36.1 | **0.001** | 0.88 | **<0.001** | 10.1 | 2.8 |
| brown | 2186 | 35.6 | **0.001** | -0.91 | **<0.001** | 24.1 | 9.5 |
| cyan | 320 | 20.3 | **0.008** | -0.61 | 0.09 | 11.5 | 4.5 |
| darkgreen | 136 | 0.1 | 0.8 | -0.01 | 1 | 10.7 | 6.1 |
| darkgrey | 108 | 0.0 | 0.9 | -0.08 | 0.0 | 12.7 | 3.9 |
| darkorange | 94 | 16.7 | **0.01** | -0.61 | 0.09 | 6.9 | 3.4 |
| darkred | 141 | 6.5 | 0.08 | -0.56 | 0.1 | 7.5 | 2.2 |
| darkturquoise | 126 | 0.8 | 0.5 | 0.42 | 0.3 | 4.1 | 2.7 |
| green | 905 | 3.7 | 0.2 | 0.50 | 0.2 | 13.1 | 13.7 |
| greenyellow | 400 | 0.3 | 0.7 | -0.07 | 0.9 | 35.9 | 16.7 |
| grey60 | 209 | 3.3 | 0.2 | -0.58 | 0.1 | 3.8 | 8.3 |
| lightcyan | 210 | 4.6 | 0.1 | 0.28 | 0.4 | 4.4 | 5.9 |

| | | | | | | | |
|---|---|---|---|---|---|---|---|
| lightgreen | 195 | 6.6 | 0.08 | -0.37 | 0.3 | 48.5 | 21.8 |
| lightyellow | 159 | 2.3 | 0.2 | 0.43 | 0.3 | 17.1 | 15.6 |
| magenta | 765 | 4.8 | 0.1 | -0.06 | 0.9 | 7.6 | 6.0 |
| midnightblue | 218 | 2.5 | 0.2 | 0.39 | 0.3 | 26.9 | 8.8 |
| orange | 95 | 0.4 | 0.6 | -0.22 | 0.6 | 2.4 | -0.3 |
| pink | 874 | 0.8 | 0.5 | 0.36 | 0.3 | 8.4 | 9.2 |
| purple | 649 | 8.0 | 0.06 | 0.63 | 0.09 | 23.9 | 6.6 |
| red | 899 | 1.3 | 0.4 | 0.34 | 0.3 | 43.5 | 9.3 |
| royalblue | 144 | 2.1 | 0.2 | 0.52 | 0.12 | 4.9 | 4.0 |
| salmon | 360 | 3.0 | 0.2 | 0.27 | 0.5 | 5.9 | 1.5 |
| skyblue | 46 | 0.7 | 0.5 | -0.35 | 0.3 | 1.2 | 1.6 |
| tan | 389 | 11.7 | **0.03** | -0.73 | **<0.001** | 4.1 | 0.3 |
| turquoise | 3880 | 3.1 | 0.2 | -0.43 | 0.3 | 71.6 | 48.0 |
| white | 93 | 5.5 | 0.1 | 0.40 | 0.3 | 1.6 | 0.7 |
| yellow | 1191 | 4.5 | 0.1 | -0.37 | 0.3 | 25.0 | 28.8 |

**Table S8.** Linkage Map anchored reference genome. Results from AllMaps-based anchoring and orienting of the reference genome scaffolds.

| | Anchored | Oriented | Unplaced |
|---|---|---|---|
| Markers (unique) | 9,297 | 9,130 | 40 |
| Markers per Mb | 20.1 | 19.9 | 0.3 |
| N50 Scaffolds | 15 | 15 | 0 |
| Scaffolds | 186 | 43 | 42,034 |
| Scaffolds with 1 marker | 145 | 14 | 34 |
| Scaffolds with 2 markers | 11 | 3 | 3 |
| Scaffolds with 3 markers | 1 | 0 | 0 |
| Scaffolds with >=4 markers | 29 | 26 | 0 |
| Total bases | 462,836,944 (74.6%) | 458,545,639 (74.0%) | 157,199,691 (25.4%) |

**Supplementary file 1:** XLSX file with GO enrichment results.
